# Graft-Induced Reprogramming Under Arterial Hemodynamic Stress is Associated with a Shared Proteomic Phenotype in Arterial and Venous Conduits Following Coronary Artery Bypass Grafting

**DOI:** 10.64898/2026.01.04.697589

**Authors:** Eun Na Kim, Suk Ho Sohn, Jiyoung Yu, Joon Seo Lim, Jiwon Koh, Jaemoon Koh, Changha Hwang, Kyunggon Kim, Ho Young Hwang, Se Jin Oh

## Abstract

**Background:** Autologous venous conduits remain the mainstay of coronary artery bypass grafting (CABG); however, their long-term patency is limited by adverse vascular remodeling. While arterial and venous grafts undergo structural and molecular adaptation to arterial hemodynamics, the mechanisms underlying successful adaptation remain poorly characterized owing to the scarcity of functioning human graft tissues.

**Methods:** We analyzed bypass grafts explanted en bloc during orthotopic heart transplantation, comprising patent vein grafts (VGP, n=19), occluded vein grafts (VGO, n=9), and patent arterial grafts (AGP, n=13), alongside freshly harvested controls (vein, n=23; artery, n=11). Quantitative histomorphometry, label-free LC–MS/MS proteomics, pathway analysis, and immunohistochemistry were performed.

**Results:** While patent arterial and venous grafts exhibited medial elastin reinforcement, adventitial neovascularization, and organized intimal remodeling, occluded vein grafts demonstrated elastic layer disruption, medial degeneration, and collagen-dense fibrosis. Proteomic analysis revealed a convergent adaptive signature in patent grafts enriched for RNA metabolism, protein synthesis, cytoskeletal dynamics, and extracellular matrix organization, with shared activation of NR4A3, STAT1, and EGFR signaling. Vein-specific adaptation involved SRC–PTGES activation with concomitant STAT3 suppression, whereas arterial grafts engaged IGF1–RUNX2-associated proliferative programs. Failed venous grafts exhibited angiotensin-driven fibroinflammatory remodeling. Endothelial nitric oxide synthase expression was markedly upregulated in the luminal endothelium and adventitial neovessels of patent grafts.

**Conclusions:** Despite distinct vascular origins, arterial and venous grafts converged toward a shared adaptive phenotype characterized by structural maturation and vasoprotective signaling. Targeting common adaptive pathways—including NR4A3, STAT1, EGFR, and adventitial eNOS—may offer therapeutic strategies to enhance long-term graft durability.

**WHAT ARE THE CLINICAL IMPLICATIONS?:** This study provides human tissue–based evidence that patent coronary bypass grafts share a common adaptive remodeling program, irrespective of whether the conduit is arterial or venous. Rather than simple venous “arterialization,” successful grafts converged on a phenotype characterized by coordinated biosynthetic activity, structural maturation, and vasoprotective signaling, whereas occluded vein grafts exhibited a distinct angiotensin-driven fibroinflammatory remodeling program. Importantly, patent grafts demonstrated conserved upregulation of NR4A3, STAT1, EGFR, and endothelial nitric oxide synthase (eNOS) in both luminal and adventitial endothelial compartments, identifying molecular features that distinguish adaptive from maladaptive remodeling in humans.

These findings are clinically relevant because they highlight shared, potentially targetable pathways that may support graft durability across conduit types and provide mechanistic support for surgical strategies that preserve perivascular adipose tissue during vein harvesting. Moreover, the association of graft failure with angiotensin-centered extracellular matrix remodeling suggests a modifiable biological axis that warrants further investigation. Together, this work establishes a human molecular framework for graft adaptation that may inform future biomarker development, therapeutic targeting, and precision strategies aimed at improving outcomes after coronary artery bypass grafting.

## INTRODUCTION

Coronary artery bypass grafting (CABG) remains a primary revascularization strategy for multivessel coronary artery disease; however, patient outcomes continue to heavily depend on long-term graft performance^1^. Arterial grafts, characterized by their elastic architecture and intrinsic compatibility with the high-pressure coronary circulation, exhibit superior long-term patency^2^. In contrast, venous grafts are abruptly exposed to arterial hemodynamics that differ fundamentally from their low-pressure venous origin and must undergo extensive adaptive remodeling to maintain patency^3^. Although arterial conduits—including the internal mammary artery, radial artery, and other elastic or muscular arteries—provide the most durable options^4^, their limited availability and anatomical constraints often necessitate the use of venous grafts in patients with multivessel disease^5^. Consequently, mitigating venous graft failure has become a key determinant of long-term outcomes following CABG.

Failure of this adaptive process—particularly in venous grafts—precipitates early thrombosis, excessive intimal hyperplasia, and accelerated atherosclerosis, events that collectively contribute to their significantly lower patency compared to arterial conduits^6,7^. Because graft failure can result in recurrent ischemia, repeat revascularization, and even sudden cardiac death, elucidating the biological mechanisms that preserve graft integrity is essential for improving CABG outcomes.

Although the mechanisms contributing to venous graft failure have been extensively described, the biological processes that enable successful graft adaptation remain poorly understood. Previous human studies have primarily examined conduits at the time of harvesting^8^ or on grafts removed following clinical failure^9^, offering limited insight into the structural and molecular changes that occur in vivo during graft function. As a result, the histomorphological features, molecular pathways, and vasoprotective mechanisms that support adaptation to coronary hemodynamic stress remain incompletely characterized.

Recent attention has focused on strategies such as no-touch harvesting and the preservation of perivascular adipose tissue (PVAT), both of which have been associated with improved graft performance^10,11^. Proposed mechanisms include enhanced endothelial integrity and increased endothelial nitric oxide synthase (eNOS) expression in freshly harvested no-touch conduits^12,13^. However, these observations are limited to pre-implant tissue, and direct evidence linking PVAT-associated or endothelial eNOS to graft patency in humans remains lacking.

A further conceptual gap concerns the traditional assumption that venous conduits undergo “arterialization” after implantation^3^. Although this term is widely used to describe the adaptive transition of veins exposed to arterial pressure, the molecular basis, temporal progression, and histomorphological correlates of this process in clinically patent grafts have never been systematically defined. It remains uncertain whether venous conduits undergo a unidirectional transition toward an arterial phenotype while arterial grafts largely preserve their native architecture, or whether more complex adaptive programs are involved.

In this study, we leveraged a rare clinical circumstance that offers uniquely valuable biological insights by enabling the direct examination of both arterial and venous bypass grafts explanted en bloc at the time of orthotopic heart transplantation. Using an integrated approach that combines quantitative histomorphology, high-resolution proteomics, and targeted immunohistochemistry, we examined not only occluded venous grafts but also successfully functioning venous and arterial grafts. By contrasting these specimens, we aimed to define the adaptive remodeling pathways, signaling trajectories, and conduit-specific mechanisms that support graft patency in humans.

## METHODS

### Study Approval and Patient Samples

This study was approved by the institutional review boards of Seoul National University Hospital (J-2306-002-1435) and SMG-SNU Boramae Medical Center (10-2023-25) on June 2, 2023. Written informed consent was obtained from all participants.

A total of eight patients who had previously undergone CABG and subsequently received orthotopic heart transplantation for end-stage heart failure (Figure 1A) were enrolled. Patient demographics are summarized in Table S1. All vein grafts were saphenous vein conduits, and all artery grafts were internal mammary arteries. Accordingly, the terms “vein graft” and “artery graft” were used to refer to these respective conduit types.

**Figure 1.**
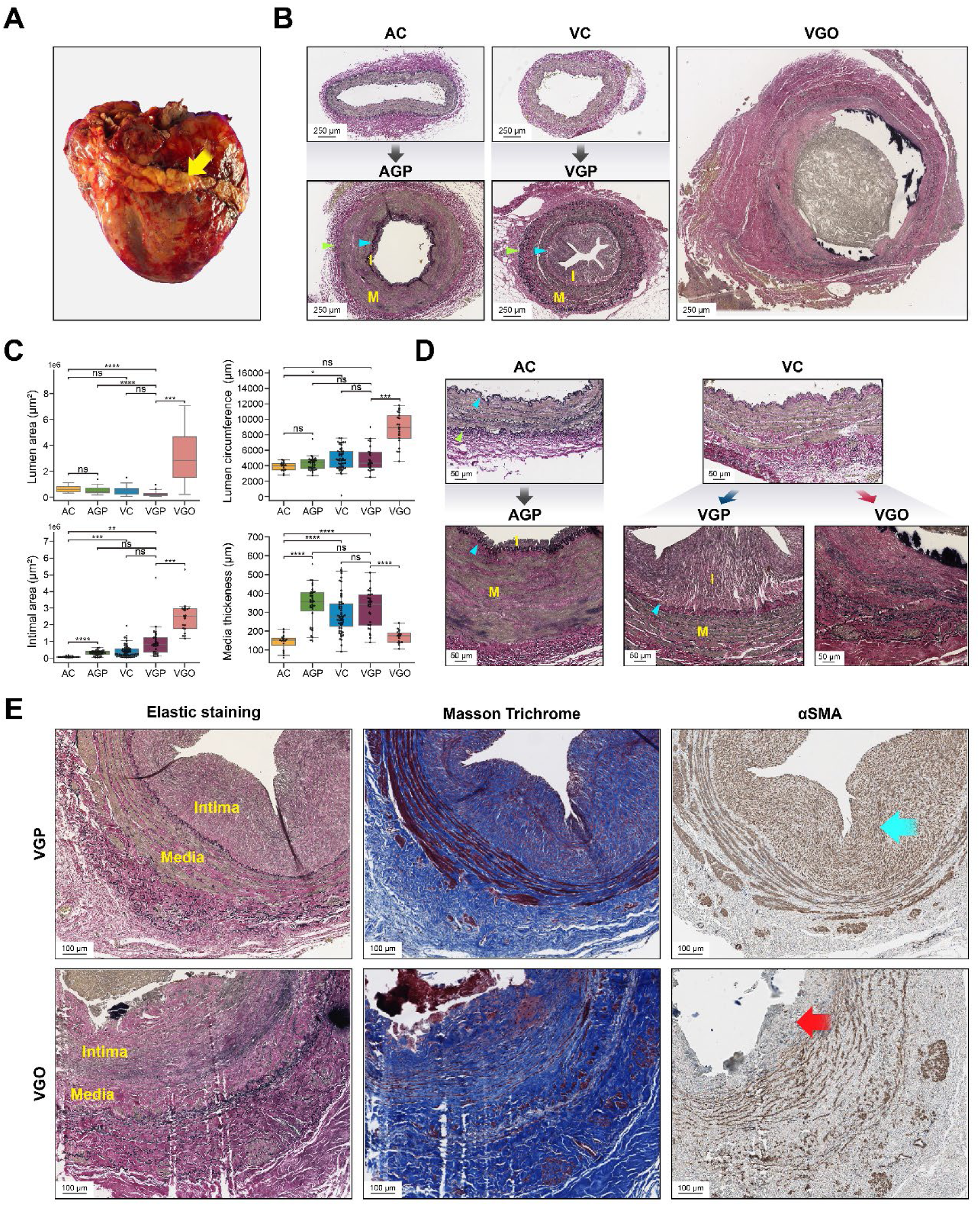
(A) Explanted heart with attached saphenous vein grafts harvested using the no-touch technique. Gross photograph of an explanted heart from a patient with prior coronary artery bypass grafting (yellow arrow). The saphenous vein graft conduits remain attached to the heart, with perivascular supportive tissue preserved as a result of the no-touch harvesting technique. (B) Histomorphologic remodeling of arterial and venous conduits following grafting. Blue arrowheads indicate the internal elastic lamina, marking the boundary between the intima (I) and media (M). Green arrowheads indicate the external elastic lamina, marking the boundary between the media (M) and adventitia. Yellow labels denote the intima (I) and media (M) within each graft. (C) Quantitative morphometric analysis of arterial and venous graft remodeling. Boxplots show the lumen area (upper left), lumen circumference (upper right), intimal area (lower left), and medial thickness (lower right) across native arterial conduits (AC), arterial grafts (AGP), native venous conduits (VC), patent venous grafts (VGP), and occluded venous grafts (VGO). Statistical comparisons are indicated above each plot (*P<0.05; **P<0.01; ***P<0.001; ****P<0.0001; ns, not significant). (D) Intimal and medial morphology across arterial and venous graft states. Elastic staining of native arteries (AC) and veins (VC), their patent graft counterparts (AGP, VGP), and occluded vein grafts (VGO). (E) Histochemical and immunohistochemical characteristics of patent versus occluded vein grafts. Elastic staining showed a well-organized intima–media structure in patent vein grafts (VGP), whereas occluded grafts (VGO) exhibited disrupted elastic layers. Masson Trichrome staining demonstrated structured fibrosis with preserved layering in VGP, but dense, disorganized collagen in VGO. αSMA staining revealed abundant, organized smooth muscle–type cells in the VGP intima (blue arrow) and near-complete absence of αSMA-positive cells in the fibrotic VGO intima (red arrow).

Seven patients underwent both procedures at Seoul National University Hospital and SMG-SNU Boramae Medical Center and received vein grafts harvested using the no-touch technique, configured in Y- or I-graft arrangements, and anastomosed to the in-situ arterial graft. One patient, who underwent CABG at an external institution, received conventionally harvested vein grafts with aortocoronary anastomosis. Graft patency prior to transplantation was assessed using coronary angiography or multi-detector computed tomography. The median interval from CABG to transplantation was 74 months (range, 0.2–226 months).

### Histomorphological Analysis

For histological assessment, we analyzed 41 formalin-fixed, paraffin-embedded (FFPE) graft blocks, including 13 artery graft patent (AGP) blocks from 5 patients, 19 vein graft patent (VGP) blocks from 6 patients, and 9 vein graft obstructed (VGO) blocks from 2 patients. As controls, freshly harvested graft tissues were obtained from 28 patients undergoing CABG, yielding 11 artery control (AC) blocks from 11 patients and 23 vein control (VC) blocks from 23 patients.

FFPE tissue blocks were sectioned into 4-μm-thick slides and stained with hematoxylin and eosin, elastin, and Masson’s trichrome (MT) to evaluate histological features, including elastic fiber content, the formation of internal and external elastic laminae, and intimal fibrosis. Slides were digitized using a whole-slide scanner (Aperio GT450 DX, Leica Biosystems). Manual annotations were performed using QuPath (https://qupath.github.io/), followed by quantification of intima, media, and adventitia thickness and area, as well as luminal, medial, and adventitial circumferences. Additionally, the number and depth of clefts within the hyperplastic intima were quantified (Figure S1).

Details on the methodologies regarding proteomic analysis, LC-MS/MS, database search, canonical pathway analysis, immunohistochemistry, and statistical analysis are provided in **Supplemental Methods**.

## RESULTS

### Histomorphological Characteristics of Patent Arterial and Venous Grafts

We analyzed 41 FFPE vessel and graft specimens to characterize baseline vascular architecture and graft-associated morphological remodeling (Figure 1B-D). After exposure to arterial hemodynamics, VGP underwent pronounced structural remodeling, characterized by substantial intimal and medial thickening, the development of a well-defined internal elastic lamina, and an increased presence of elastic fibers interwoven within the medial layer. In addition, a newly formed, wavy elastic layer between the media and adventitia was consistently observed, representing a structural reorganization that renders the venous wall similar to the layered organization typical of arterial conduits. In contrast, VGO showed aneurysmal dilatation, loss of organized elastic layers, medial degeneration, and collapse of the normal intima–media–adventitia boundaries, consistent with structural failure rather than adaptive remodeling. Detailed results of quantitative morphometric analysis and intimal measurements are provided in **Supplemental Results**.

### Immunohistochemical Analysis of Patent and Occluded Vein Grafts

Histochemical and immunohistochemical staining (Figure 1E) revealed distinct differences in intimal organization between patent and occluded vein grafts. In VGP, elastic staining demonstrated a well-organized intima and media. Masson’s trichrome staining indicated fibrosis; however, the collagen matrix maintained a structured pattern with interspersed αSMA-positive myofibroblasts, suggesting an orderly and adaptive remodeling process. In contrast, VGO displayed a fundamentally different morphology. Elastic staining revealed significant disruption of the elastic lamina structures. Masson’s Trichrome staining demonstrated dense, disorganized collagen consistent with a non-cellular fibrotic matrix, and αSMA staining confirmed the absence of smooth muscle–type cells within the thickened intima.

### Proteomics

#### Global Proteomic Profiling Reveals Condition-Specific Molecular Signatures

To characterize the molecular landscape of vascular remodeling during graft adaptation and failure, we performed quantitative proteomic analysis using LC-MS/MS on five experimental groups: VC, AC, VGP, AGP, and VGO. A total of 4,836 proteins were identified across all vascular samples (Figure 2A). Unsupervised two-dimensional PCA revealed a clear separation between AC and VC samples. In contrast, AGP and VGP samples clustered closely together. VGO samples formed a distinct, isolated cluster, separate from both native vessels and patent grafts (Figure 2B).

**Figure 2.**
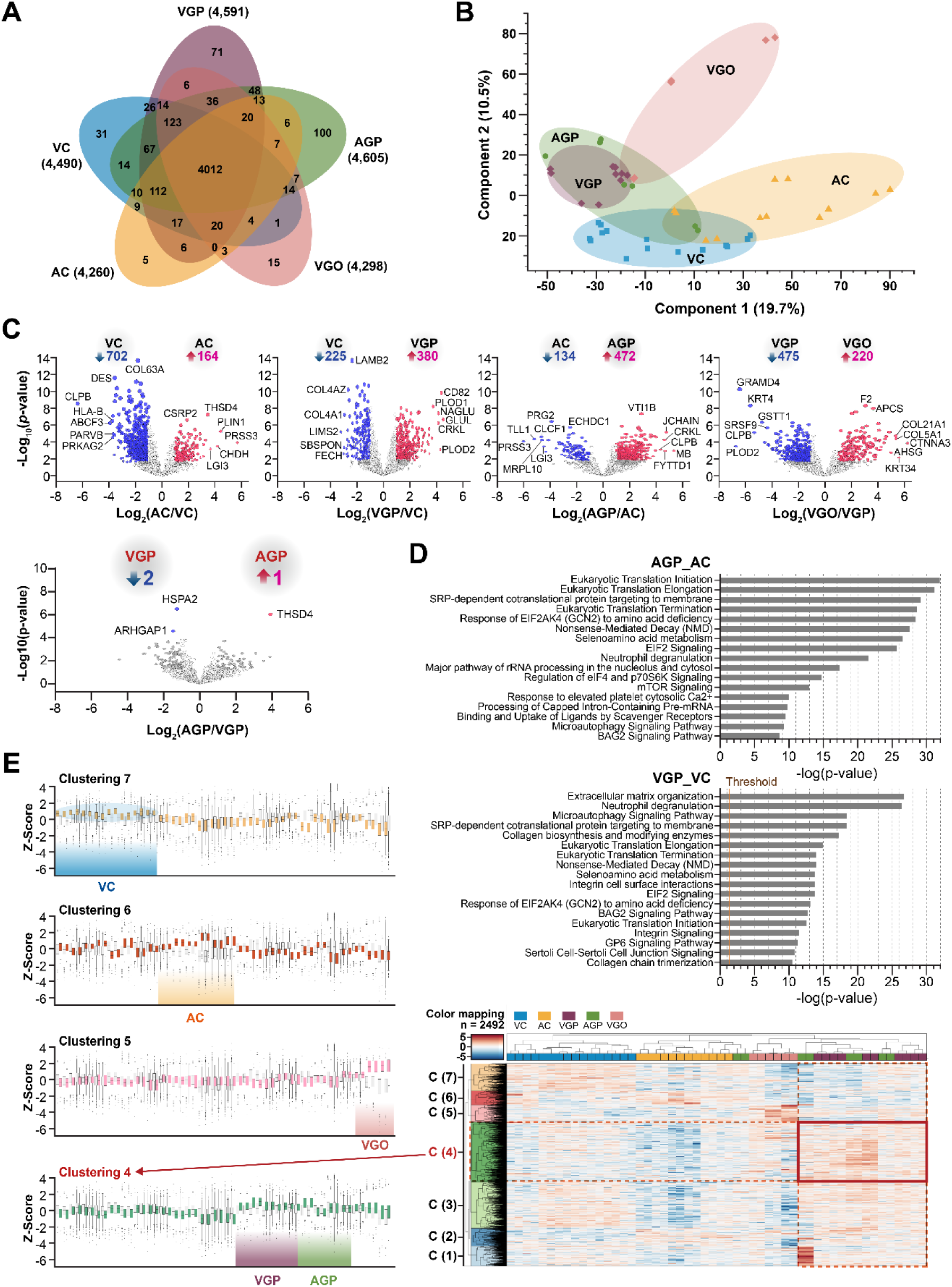
(A) Number of identified proteins Venn diagram showing overlap of identified proteins across five vascular tissue groups. A total of 4,836 proteins were identified across all samples using mass spectrometry–based proteomic analysis. The five groups included vein control (VC, n=4,490 proteins), artery control (AC, n=4,260), patent vein grafts (VGP, n=4,591), patent artery grafts (AGP, n=4,605), and occluded vein grafts (VGO, n=4,298). Among these, 4,012 proteins were commonly detected across all five groups, while subsets of proteins were uniquely identified in specific conditions. The number of shared and group-specific proteins is indicated within each section of the Venn diagram. (B) Unsupervised 2-dimensional PCA showing distinct clustering patterns across vascular groups. AC and VC samples separated widely along PC1, reflecting their differing baseline proteomic profiles. AGP and VGP samples clustered closely, indicating similar post-grafting proteomic changes. VGO samples formed a cluster that was distinct from both native vessels and patent grafts. (C) Differential protein expression among native vessels and graft types. Volcano plot comparing VC and AC showing 702 proteins enriched in VC and 164 enriched in AC, indicating clear baseline proteomic differences between veins and arteries. Comparison of AC and AGP identified 134 proteins enriched in AC and 472 enriched in AGP, consistent with extensive arterial graft remodeling. Comparison of VC and VGP revealed 225 proteins enriched in VC and 380 enriched in VGP, reflecting substantial remodeling following venous grafting. Direct comparison of AGP and VGP showed only a minimal number of differentially expressed proteins, underscoring the post-grafting similarity between arterial and venous conduits. Comparison of VGP and VGO identified 475 proteins enriched in VGP and 220 enriched in VGO, highlighting the molecular divergence between functional and occluded vein grafts. (D) Canonical pathway enrichment analysis of arterial and venous grafts. (Left) Canonical pathway enrichment analysis for AGP versus AC showed significant activation of translation-related pathways (e.g., initiation, elongation, termination, and SRP-dependent targeting), EIF2/GCN2 signaling, nonsense-mediated decay, and proteostasis-associated programs, including microautophagy and BAG2 signaling. (Right) Canonical pathway enrichment analysis for VGP versus VC showed similar enrichment in translation-associated pathways, including EIF2/GCN2, and proteostasis pathways, along with additional processes related to extracellular matrix organization, collagen biosynthesis, integrin signaling, and neutrophil degranulation. (E) Hierarchical clustering reveals group-specific proteomic signatures across vascular samples. Clustering of the 2,492 most variable proteins generated distinct groups corresponding to major vascular phenotypes. Cluster 7 consisted predominantly of VC samples, while Cluster 6 was largely composed of AC samples, reflecting baseline proteomic differences between native veins and arteries. Cluster 5 was enriched with VGO samples and displayed a molecular pattern distinct from all other groups. Cluster 4 contained proteins consistently upregulated in both AGP and VGP, highlighting the close similarity of proteomic profiles in functional arterial and venous grafts.

#### Volcano Plot Analysis of Graft-Associated Proteomic Changes

Volcano plot analysis (Figure 2C) revealed extensive proteomic remodeling across all graft types. Only a very small subset of proteins differed between AGP and VGP, underscoring the proteomic similarity between the two conduits after grafting. In contrast, the comparison between VGP and VGO showed marked divergence, with 475 proteins enriched in VGP and 220 enriched in VGO.

#### Canonical Pathway Enrichment Analysis of AGP and VGP

Canonical pathway enrichment analysis (Figure 2D) demonstrated that AGP (vs. AC) and VGP (vs. VC) share a broad set of commonly enriched pathways. Both comparisons showed significant activation of translation-associated programs, indicating similarities in biosynthetic and proteostasis-related pathways in both graft types.

#### Hierarchical Clustering Revealed Group-Specific Proteomic Signatures

Hierarchical clustering of the 2,492 differentially expressed proteins revealed distinct group-specific expression patterns across vascular samples (Figure 2E)—VC in Cluster 7, AC in Cluster 6, VGO in Cluster 5, and both AGP and VGP in Cluster 4. To further characterize the biological processes represented by each cluster, we employed a three-tiered analytical workflow consisting of: (1) pathway enrichment analysis using Ingenuity Pathway Analysis, which offers a function-centered interpretation of overrepresented biological programs; (2) protein–protein interaction (PPI) network analysis using the STRING database^14^, which delineates the structural and functional connectivity among proteins; and (3) integrative functional annotation using the Metascape^15^ enrichment bubble plot (GO/KEGG/Reactome). These analyses were conducted on cluster-specific protein subsets derived from the hierarchical clustering: AC (Cluster 6, n=171 proteins), VC (Cluster 7, n=340 proteins), VGP + AGP (Cluster 4, n=741 proteins), and VGO (Cluster 5, n=210 proteins).

#### Integrated Functional Analysis Revealed Fundamentally Distinct Remodeling Programs Represented by Clusters 4 and 5

Cluster 4 (VGP + AGP) captured the shared proteomic signature of patent arterial and venous grafts. Pathway enrichment analysis (Figure 3A) demonstrated coordinated activation of RNA metabolism, mRNA processing, translation initiation and elongation, ribonucleoprotein biogenesis, and SRP-dependent cotranslational protein targeting to membranes, along with pathways supporting extracellular matrix (ECM) organization and cytoskeletal adaptation.

**Figure 3.**
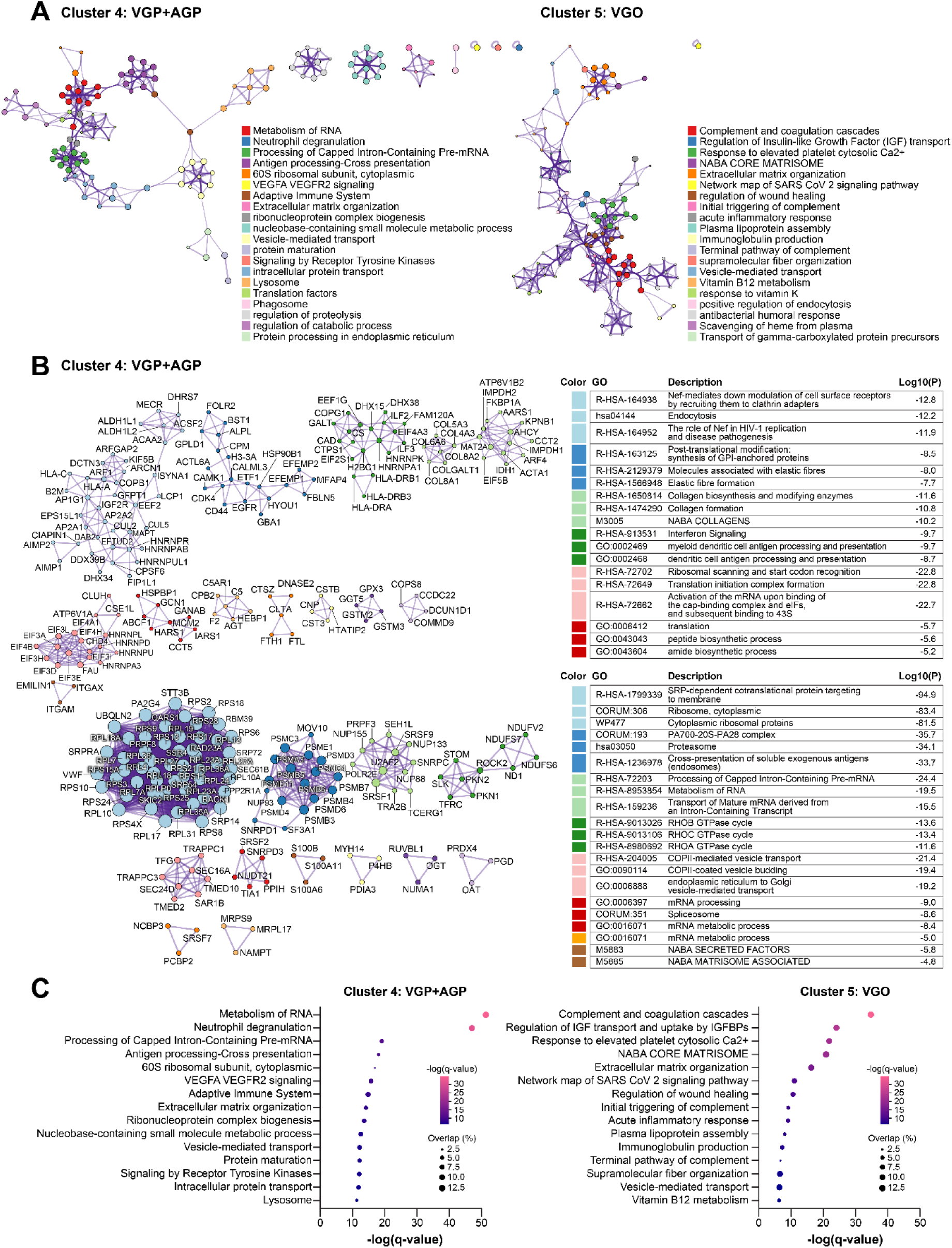
Integrated functional analysis differentiates adaptive remodeling in patent grafts from maladaptive processes in occluded grafts. (A) Pathway enrichment analysis of Cluster 4 (VGP + AGP) and Cluster 5 (VGO). Cluster 4 showed coordinated enrichment in RNA metabolism, mRNA processing, translation initiation/elongation, ribonucleoprotein biogenesis, and pathways supporting extracellular matrix (ECM) organization and cytoskeletal adaptation. In contrast, Cluster 5 exhibited predominant enrichment of complement and coagulation cascades, acute inflammatory signaling, wound healing, and disorganized ECM remodeling. (B) Protein–protein interaction (PPI) networks for Cluster 4 Cluster 4 networks revealed densely interconnected ribosomal/translation hubs and organized ECM-remodeling subnetworks, consistent with an active, reparative remodeling phenotype. (C) Metascape pathway enrichment analysis of functional and obstructed grafts (Cluster 4 and Cluster 5). Pathway enrichment analysis was conducted using Metascape (www.metascape.org), which integrates multiple curated ontologies, including GO, KEGG, and Reactome. (A) Pathways enriched in functional grafts, including vein graft patent (VGP) and artery graft patent (AGP) groups. (B) Pathways enriched in the vein graft obstructed (VGO) group. Each bubble represents a biological pathway, where the x-axis shows enrichment significance (–log₁₀(q-value)), the y-axis shows pathway names, bubble size reflects the percentage of overlapping genes (Overlap%), and bubble color reflects significance strength, where a more intense pink indicates higher significance. Cluster 4 (VGP + AGP) demonstrated enrichment of pathways related to RNA metabolism, VEGFA–VEGFR2 signaling, receptor tyrosine kinase activity, protein maturation, intracellular transport, lysosomal processes, and immune-regulatory programs—reflecting a biosynthetically active, structurally adaptive, and immune-balanced phenotype characteristic of functional grafts. In contrast, Cluster 5 (VGO) was enriched for complement and coagulation pathways, NABA matrisome and ECM-associated programs, platelet activation, plasma lipoprotein assembly, and IGFBP-mediated IGF transport regulation—defining a matrix-disruptive, thrombosis-prone, and growth factor–dysregulated environment associated with graft obstruction.

PPI network analysis (Figure 3B) revealed densely interconnected ribosomal and translation hubs, as well as organized ECM-remodeling subnetworks, consistent with an active, reparative remodeling phenotype. In contrast, Cluster 5 (VGO) represented the molecular landscape of failed vein grafts. Pathway enrichment analysis revealed predominant activation of complement and coagulation cascades, acute inflammatory signaling, wound healing, and disorganized ECM remodeling. PPI networks (Figure S4) formed extensive thrombo-inflammatory modules and collagen-rich ECM clusters, indicating a shift toward a maladaptive, pro-inflammatory, and pro-thrombotic phenotype.

Metascape-based functional annotation (Figure 3C) further distinguished the adaptive and maladaptive programs of these clusters. Cluster 4 showed enrichment in RNA metabolism, VEGFA–VEGFR2 signaling, receptor tyrosine kinase pathways, protein maturation, intracellular transport, lysosomal activity, and immune-regulatory processes, reflecting a biosynthetically vigorous, structurally adaptive, and immunologically balanced program characteristic of successful grafts. In contrast, Cluster 5 was enriched in complement and coagulation cascades, NABA matrisome pathways, extracellular matrix organization, platelet activation, plasma lipoprotein assembly, and IGFBP-mediated IGF transport regulation. This enrichment is consistent with a matrix-disruptive, thrombosis-prone, and growth factor–dysregulated environment associated with graft obstruction.

Canonical pathway analysis further revealed distinct activation patterns across graft conditions, with biosynthetic and vascular remodeling pathways predominating in functional grafts, while inflammatory and coagulative pathways characterized occluded grafts.

#### Activation of Proliferative and Biosynthetic Programs and Suppression of Contractile Regulators in Patent Grafts

Upstream regulator analysis of patent grafts compared to native vessels revealed coordinated activation of proliferative and biosynthetic signaling networks. In AGP, key activated upstream nodes included IGF1, RUNX2, NR4A3, PGC-1α, EGFR, and MYC (Figure 4A). In VGP, STAT1, SRC, PTGES, resistin (RETN), and G6PD were identified as activated nodes, forming a coherent regulatory network (Figure 4A,B). In contrast, regulators involved in vascular contractility and cytoskeletal integrity, including serum response factor (SRF) and myocardin-related transcription factor A (MRTFA), were predicted to be inhibited in VGP. This transcriptional profile indicates phenotypic modulation toward a synthetic phenotype, characterized by enhanced extracellular matrix production, increased biosynthetic capacity, and elevated proliferative activity—hallmarks of adaptive vascular remodeling during vein graft transformation in the arterial circulation.

**Figure 4.**
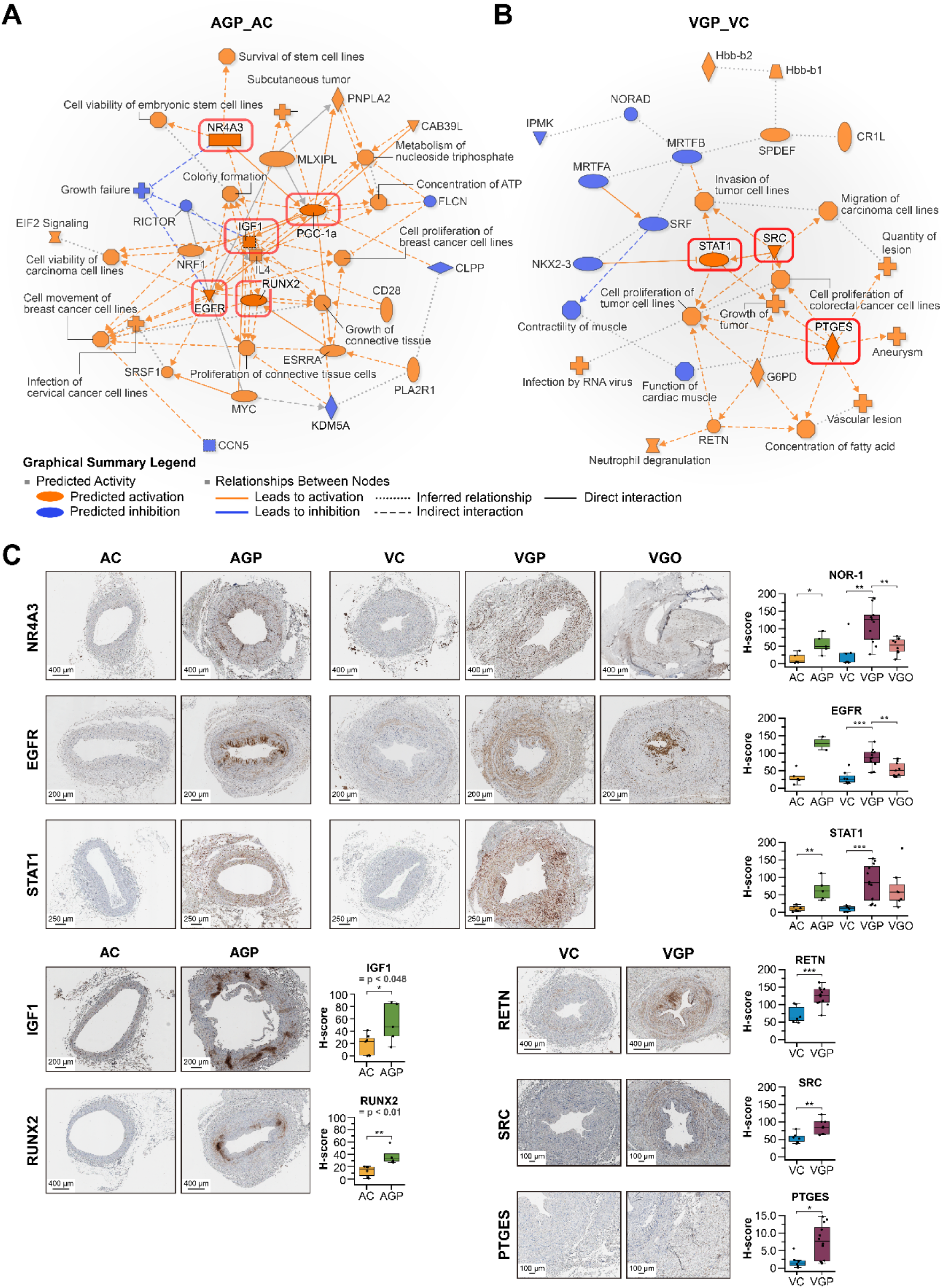
Upstream regulatory programs and immunohistochemical validation in arterial and venous grafts (A) Arterial grafts (AGP vs. AC). Upstream regulator analysis revealed coordinated activation of metabolic and proliferative regulators, including IGF1, RUNX2, NR4A3, PGC-1α, EGFR, and MYC. (B) Venous grafts (VGP vs. VC). Patent vein grafts showed activation of proliferative–biosynthetic regulators, including STAT1, SRC, PTGES, RETN, and G6PD, accompanied by predicted inhibition of vascular contractility regulators such as SRF and MRTFA—indicating a shift toward a synthetic, remodeling-competent phenotype. (C) Immunohistochemical validation. IHC confirmed increased expression of NR4A3, EGFR, and STAT1 in both AGP and VGP (p<0.01), identifying these as conserved adaptive regulators across graft types. Arterial grafts showed specific upregulation of IGF1 and RUNX2, while venous grafts exhibited higher expression of RETN, SRC, and PTGES compared with their respective controls, validating graft-type–specific adaptive signaling predicted by upstream analyses.

#### Immunohistochemical Validation of Key Regulatory Nodes

To validate pathway-level predictions, immunohistochemical analysis was performed on key upstream regulators identified through IPA analysis (Figure 4C). Shared adaptive regulators across both graft types—NR4A3, EGFR, and STAT1—demonstrated significantly increased expression in both arterial grafts (AGP vs. AC, P<0.01) and venous grafts (VGP vs. VC, P<0.01), establishing these as conserved adaptive regulators during adaptation to arterial hemodynamics. Notably, the expressions of NR4A3 and EGFR were markedly reduced in VGO, consistent with regression of proliferative signaling during graft failure. Graft-Type–Specific Adaptive Signatures AGP exhibited significant increases in IGF1 and RUNX2 expression compared to native AC, validating pathway predictions of growth factor–driven proliferative and osteogenic signaling in arterial graft adaptation. Conversely, VGP showed elevated expression of RETN (resistin), SRC, and PTGES compared to native VC, confirming graft-type–specific activation of metabolic stress and biosynthetic mediator pathways predicted by IPA analysis.

PGC-1α—a factor predicted by the proteomic model to vary by graft type—showed an upward trend in both arterial and venous grafts (AGP > AC; VGP > VC), although the differences did not reach statistical significance (AC vs. AGP, P=0.177; VC vs. VGP, P=0.104). Similarly, c-Myc expression tended to be higher in AGP compared with AC, but this difference was not statistically significant (P=0.114) (Figure S14).

Collectively, these immunohistochemical findings validate the pathway analysis predictions and establish both conserved adaptive programs (NR4A3, EGFR, STAT1, PGC-1α) and graft-type–specific regulatory signatures (IGF1/RUNX2 in arterial grafts; RETN/SRC/PTGES in venous grafts) that operate during vascular graft adaptation to arterial circulation.

#### EGFR–STAT3 Signaling Differences Between VGP and VGO

Upstream regulator analysis predicted preferential activation of EGFR in VGP, identifying EGFR as a key regulatory node with predicted inhibitory effects on STAT3 (Figure 5A). This prediction aligns with prior immunohistochemical findings (Figure 4C), which demonstrate elevated EGFR expression in VGP and AGP, whereas VGO exhibits marked EGFR downregulation.

**Figure 5.**
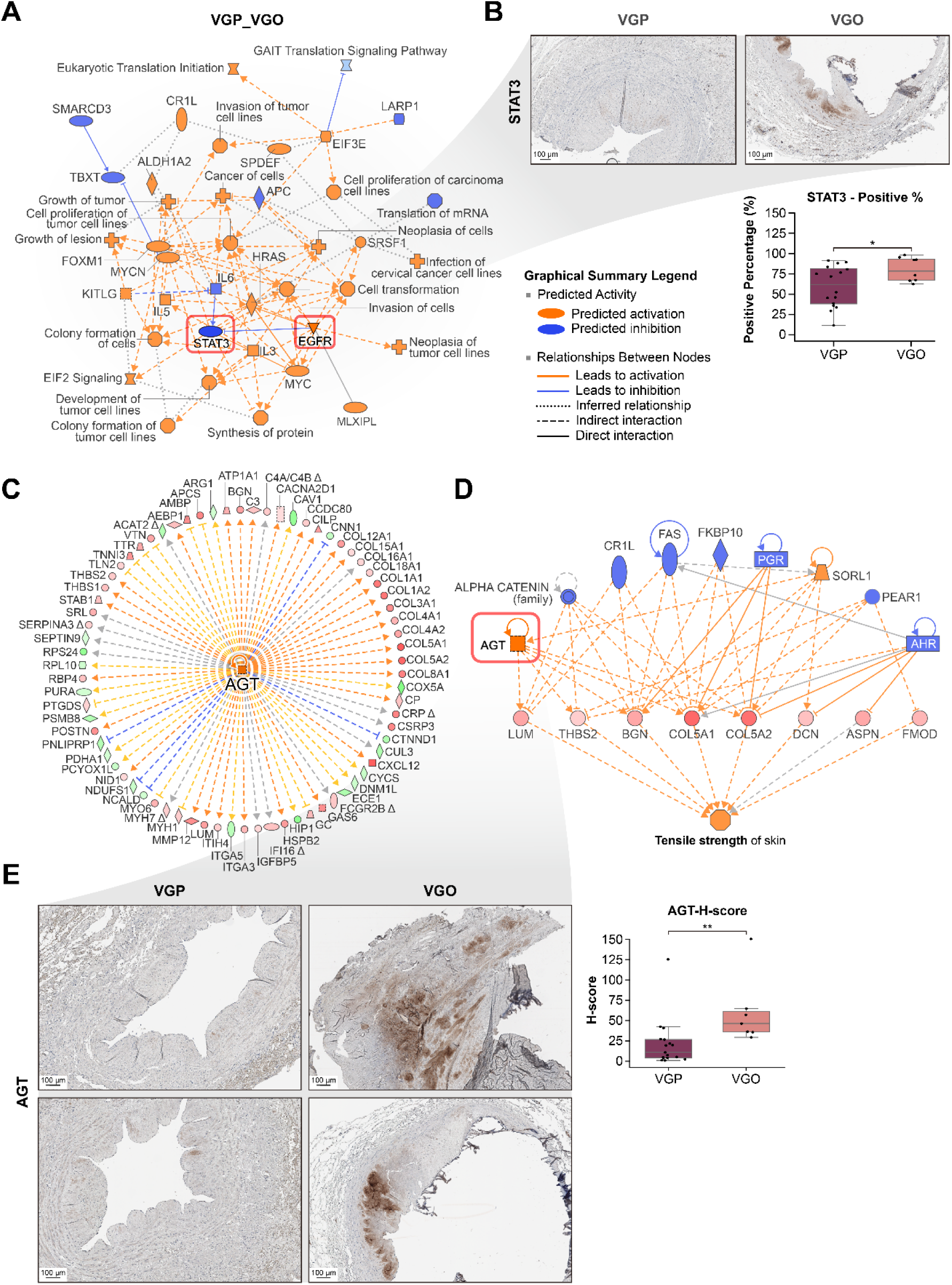
Distinct upstream signaling patterns and angiotensin-associated extracellular matrix remodeling in patent and occluded venous grafts (A) Upstream regulator analysis comparing VGP and VGO using Ingenuity Pathway Analysis. Orange nodes indicate predicted activated regulators, while blue nodes indicate inhibited regulators. In VGP relative to VGO, STAT3 downregulation, coupled with reciprocal EGFR upregulation, emerged as a key regulatory axis distinguishing successful graft adaptation. (B) STAT3 expression assessed by immunohistochemistry and quantitative analysis. Upper panels: Representative STAT3 staining in VGP (left) and VGO (right) demonstrating minimal STAT3 expression in patent grafts versus increased nuclear positivity in occluded grafts. Scale bars=100 µm. Lower panel: Quantification of STAT3-positive cells showing a trend toward elevated expression in VGO (P=0.08). n=8–10 per group. (C) Upstream regulator analysis identified angiotensin as a central activated node in VGO. The angiotensin-centered regulatory network illustrates predicted downstream targets related to the extracellular matrix. Node size reflects network connectivity, while connecting edges represent predicted regulatory or protein–protein interactions. (D) Expanded angiotensin regulatory network highlighting extracellular matrix structural proteins (orange), matrix-associated signaling molecules (blue), and downstream predicted biological functions (pink). Color intensity reflects the predicted activation state. Upregulated matrix components include fibrillar collagens (COL5A1, COL5A2), small leucine-rich proteoglycans (decorin, biglycan, lumican), and thrombospondin-2, which collectively contribute to enhanced matrix tensile strength in occluded grafts. (E) Immunohistochemical validation of the expression of angiotensinogen, the precursor of angiotensin. Left column: Two representative cases of patent venous grafts (VGP), each from a different patient, demonstrating minimal angiotensinogen staining throughout the vessel wall. Right column: Two representative cases of occluded venous grafts (VGO), each from a different patient, showing prominent angiotensinogen deposition in the media and adventitia, with dense accumulation within fibrotic regions. All images are at the same magnification. Scale bars=200 µm. Right panel: Quantitative H-score analysis revealed significantly elevated angiotensinogen expression in VGO versus VGP (**P<0.01). Box plots display the median and interquartile range (n=8–12 per group).

To evaluate the corresponding STAT3 response, immunohistochemical staining was performed (Figure 5B). VGP displayed minimal STAT3 nuclear positivity, whereas VGO demonstrated pronounced STAT3 upregulation within the fibrotic neointima. Quantitative analysis revealed a trend toward higher STAT3 expression in VGO (P=0.08), supporting reciprocal signaling dynamics characterized by EGFR-dominant/STAT3-suppressed activity in patent grafts versus STAT3-dominant activation in occluded grafts.

#### Angiotensin Activation and ECM Remodeling in VGO

Upstream regulator analysis comparing VGP and VGO identified angiotensinogen as one of the most strongly activated regulatory nodes associated with graft failure (Figure 5C). This activation was accompanied by the coordinated upregulation of multiple extracellular matrix–related proteins in VGO. The angiotensin-centered regulatory network (Figure 5D) revealed elevated expression of fibrillar collagens (COL5A1, COL5A2), small leucine-rich proteoglycans (decorin, biglycan, lumican), and thrombospondin-2, which collectively promote collagen fibrillogenesis and matrix stabilization. Regulator-effects modeling predicted enhanced matrix tensile strength as a downstream consequence in VGO, consistent with maladaptive fibrotic remodeling and reduced vascular compliance.

To validate these proteomic predictions, immunohistochemical staining for angiotensinogen was performed on matched VGP and VGO sections (Figure 5E). VGP exhibited minimal angiotensinogen immunoreactivity, showing only faint background staining. In contrast, VGO demonstrated marked angiotensinogen accumulation throughout the media and adventitia, with intense deposition in regions of dense extracellular matrix expansion and structural distortion. Quantitative analysis confirmed significantly elevated angiotensinogen expression in VGO compared with VGP (P<0.01), supporting the association between angiotensin pathway activation and fibroinflammatory remodeling in failed grafts.

#### Enhanced eNOS Expression in Luminal and Adventitial Neovascular Endothelial Cells Following Arterial Grafting

Expression of eNOS was evaluated by immunohistochemical staining across all vessel groups (Figure 6A). AC and VC demonstrated weak to moderate eNOS immunoreactivity in the luminal endothelium. Both AGP and VGP showed markedly elevated eNOS expression, whereas VGO showed minimal signal, consistent with near-complete endothelial denudation.

**Figure 6.**
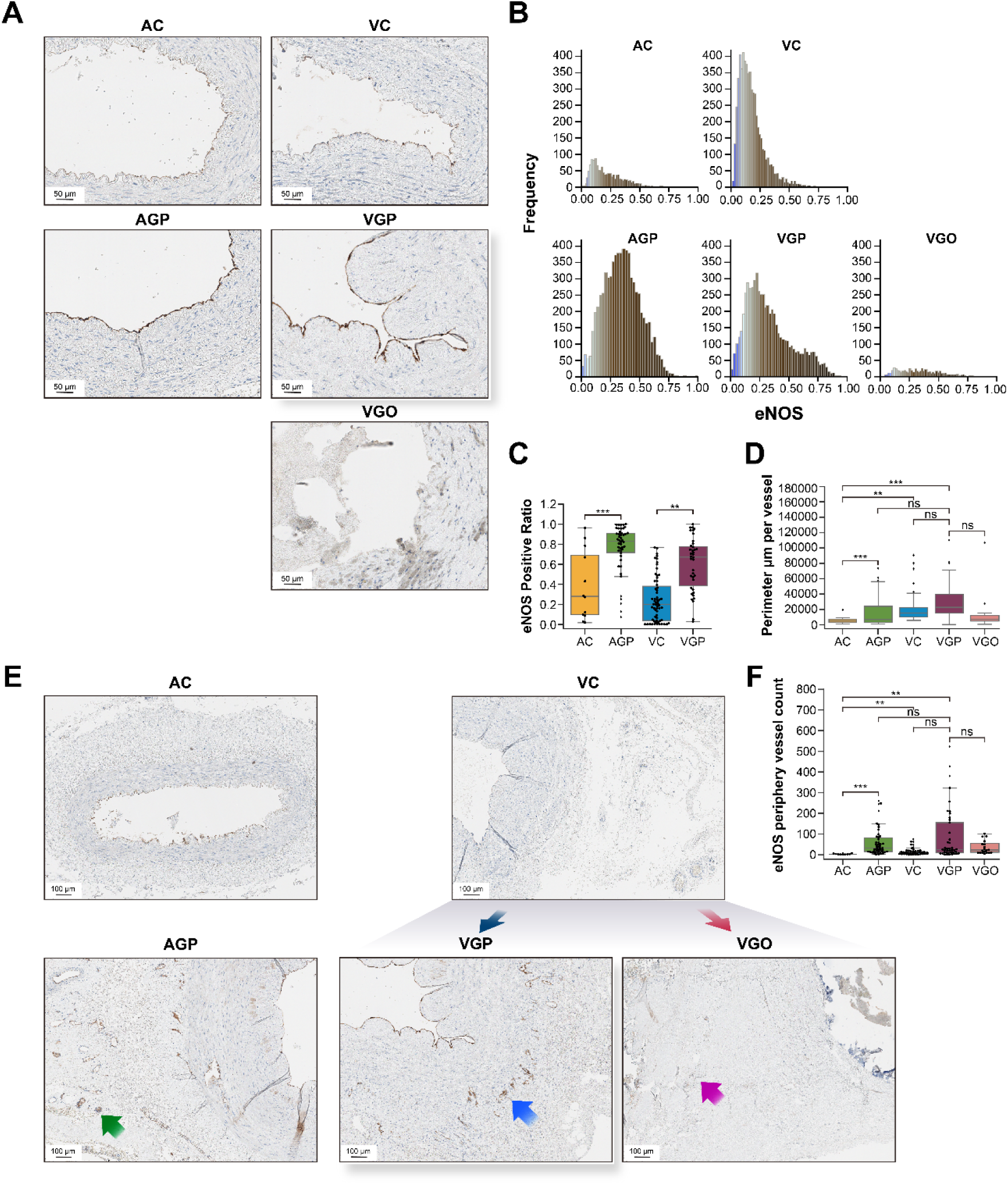
Upregulation of eNOS in luminal and adventitial endothelial cells in arterial grafts (A-C) eNOS expression in the luminal endothelium. (A) Representative immunohistochemistry showing weak eNOS expression in control vessels (AC, VC) and marked upregulation in patent grafts (AGP, VGP). Occluded grafts (VGO) exhibit endothelial denudation with minimal eNOS signal. Scale bars=50 µm. (B) Histogram analysis of single-cell optical density revealed a rightward shift in the graft endothelium, indicating enhanced eNOS expression. (C) Proportion of eNOS-positive cells (optical density >0.2) was significantly higher in AGP versus AC (***P<0.001) and in VGP versus VC (**P<0.01). (D, E) eNOS expression in the adventitial vasa vasorum. (D) Representative images show enhanced neovascularization with eNOS-positive endothelium in AGP (green arrow) and VGP (cyan arrow) compared with controls. VGO (magenta arrow) demonstrated sparse vasculature lacking eNOS expression. Scale bars=100 µm. (E) Left: Total microvascular perimeter was significantly larger in AGP versus AC (***P<0.001). Right: eNOS-positive adventitial vessel count was significantly higher in AGP versus AC (***P<0.001) and in VGP versus VC (**P<0.01). Box plots display median and interquartile range. Statistical analysis was performed using one-way ANOVA with Tukey’s post hoc test.

Single-cell optical density analysis demonstrated enhanced eNOS expression in patent grafts. Histogram distributions showed rightward shifts in AGP and VGP compared with their respective controls (Figure 6B). Using a threshold of mean DAB optical density > 0.2, the proportion of eNOS-positive cells was significantly higher in AGP versus AC (P<0.001) and in VGP versus VC (P<0.01) (Figure 6C), distinguishing successfully adapted conduits from failed grafts.

#### Adventitial Neovascularization and Associated Upregulation of eNOS

To evaluate adaptive remodeling of the vessel wall, we quantified CD31-positive adventitial neovascularization (Figure S1E–H). Total microvascular content, measured as the cumulative CD31-positive vessel perimeter, was significantly higher in AGP compared with AC (P<0.001). Venous grafts showed a trend toward increased vasa vasorum relative to VC, although this did not reach statistical significance. VGO demonstrated markedly reduced adventitial microvascular density (Figure 6D).

Notably, robust eNOS immunoreactivity was observed in the endothelial cells lining adventitial neovessels of patent grafts (Figure 6E, arrows). Quantification of eNOS-positive microvessels—defined by endothelial DAB staining with nuclear co-localization—revealed significant increases in AGP vs. AC and in VGP vs. VC (Figure 6F). VGO exhibited a marked reduction in eNOS-positive neovessels, suggesting that eNOS expression in both luminal and adventitial endothelium is associated with graft patency.

## DISCUSSION

By leveraging a unique opportunity to analyze patent arterial and venous grafts explanted en bloc during cardiac transplantation, we provide the first comprehensive molecular characterization of successful graft adaptation in humans. Our findings challenge the traditional concept of venous “arterialization” and reveal that both arterial and venous grafts converge toward a shared adaptive phenotype characterized by coordinated activation of biosynthetic, vasoprotective, and stress-responsive pathways. We also identified specific molecular signatures—including NR4A3, STAT1, EGFR, and eNOS upregulation—that distinguish patent from occluded grafts, and demonstrate that angiotensinogen-driven fibroinflammatory remodeling is associated with graft failure. These findings provide a mechanistic framework for developing targeted therapies to enhance graft durability and improve long-term outcomes following CABG.

### Histological Signatures of Graft Adaptation

While previous studies have predominantly focused on pathological processes leading to graft failure^16^, our histological analysis of functionally patent grafts reveals adaptive remodeling signatures that have been largely overlooked in the literature. Traditional conceptualizations of vein graft adaptation have emphasized “arterialization” as a unidirectional transformation toward an arterial phenotype^3^. However, our findings demonstrate that both arterial and venous grafts undergo substantial structural reorganization that does not simply recapitulate native arterial architecture.

We found that arterial grafts, rather than preserving their native elastic artery characteristics, transitioned toward a muscular artery-like configuration characterized by marked medial thickening and an augmented internal elastic lamina. In contrast, venous grafts developed features absent in native veins—including a prominent internal elastic lamina, extensive medial muscularization accompanied by elastic fiber deposition, and the formation of a novel external elastic layer. Most strikingly, despite their distinct embryological origins and baseline morphologies, both graft types converged toward similar caliber and structural organization while retaining unique architectural features not found in either native arteries or veins. This morphological convergence, coupled with the preservation of graft-specific characteristics, challenges the classical paradigm of simple arterialization and instead supports the emergence of a distinct adaptive phenotype uniquely tailored to the extraordinary hemodynamic demands of the coronary circulation. In this setting, grafts are subjected to intense pulsatile stress as the first major branches arising directly from the aortic root, receiving the full force of ventricular ejection without the dampening effect experienced by more distal vascular territories. In contrast, occluded grafts failed to achieve this adaptive remodeling, exhibiting severe aneurysmal dilatation, loss of elastic integrity, and medial necrosis — indicating that successful adaptation requires coordinated structural reorganization rather than passive exposure to arterial flow.

### Adaptive Versus Pathologic Intimal Hyperplasia in Vein Grafts

Intimal hyperplasia has traditionally been regarded as a pathological hallmark of vein graft failure in CABG, contributing to progressive luminal narrowing and eventual occlusion^17^. However, our findings fundamentally challenge this binary perspective by demonstrating that substantial intimal thickening occurs in both functionally patent and occluded grafts, albeit with markedly distinct structural and cellular phenotypes. Patent grafts exhibited well-organized intimal architecture with preserved luminal patency, suggesting that intimal hyperplasia may represent an adaptive, actively regulated biological response essential for successful graft remodeling rather than an inherently pathological process.

This reconceptualization is strongly supported by previous studies. Jiang and colleagues, using a bilateral jugular vein graft model that exposed grafts to varying hemodynamic conditions, demonstrated that intimal hyperplasia develops as an adaptive response to mechanical stress; their findings showed that grafts maintain patency through coordinated outward remodeling^18^. Similarly, a recent animal study has reported that patent grafts exhibit concurrent intimal hyperplasia alongside positive remodeling—characterized by lumen preservation despite wall thickening—indicating that intimal hyperplasia is a component of successful environmental adaptation rather than an inevitable pathological process^19^.

Histologically, patent grafts in our cohort exhibited a structurally organized intima with well-preserved endothelial integrity and a clearly demarcated internal elastic lamina, as demonstrated by elastic tissue staining. Notably, α-SMA-positive myofibroblasts were consistently identified within the intima of patent grafts, indicating controlled fibromuscular remodeling mediated by tightly regulated cellular proliferation and differentiation. The mechanistic role of myofibroblasts in adaptive intimal remodeling has been elegantly elucidated by Vasudevan and colleagues through ultrastructural analyses of rat vein grafts^20^. They documented that adventitial fibroblasts undergo coordinated migration into the intima and subsequent differentiation into myofibroblasts, which facilitate intimal thickening through organized matrix synthesis. Importantly, their ultrastructural findings revealed that adaptive intimal thickening comprises well-organized, multilayered myofibroblast assemblies rather than disorganized collagenous fibrosis—observations that closely align with our immunohistochemical characterization of human patent grafts. In contrast, occluded grafts exhibited profoundly disorganized intimal architecture, fragmented elastic laminae, and excessive extracellular matrix deposition without coordinated cellular organization. This distinction underscores that the qualitative characteristics of intimal hyperplasia—rather than merely its presence or extent—determine the long-term fate of the graft.

### Embryonic Arterialization Signaling

The convergent adaptive remodeling observed in both arterial and venous grafts may reflect the reactivation of embryonic arterialization signaling pathways. During vascular development, arterial identity is established through the coordinated activation of VEGF-Notch-ERK signaling cascades, which promote arterial-specific gene expression and structural maturation. This developmental program is particularly well-characterized in coronary artery embryogenesis, where VEGFR2-ERK-eNOS and NOTCH4 signaling axes are distinctly upregulated and activated in endothelial cells during vascular morphogenesis^21–23^. Specifically, VEGFR2 signaling activates the downstream ERK pathway, which is critical for endothelial proliferation, branching morphogenesis, and lumen formation in both coronary and general vascular embryology. eNOS is also upregulated and activated downstream of VEGFR2 in response to VEGF stimulation in the endothelium, mediating vasodilation and angiogenic responses required for vessel remodeling. Regarding NOTCH4, this receptor (along with NOTCH1) is specifically expressed in embryonic vascular endothelium and becomes markedly upregulated and activated during arterial specification and network patterning. Activation of NOTCH4 in endothelial cells is essential for correct branching morphogenesis and vascular patterning in the developing embryo, including the coronary vasculature, with increased NOTCH4 expression further supporting arterial specification from progenitor cells^22,23^

Our proteomic analysis revealed that patent grafts exhibited significant enrichment of key components of the developmental arterialization program: VEGFR2 signaling (Figure 3A and Figure S9B), ERK1/2 MAPK pathways (FDR<5×10⁻⁴; Supplementary Data), and NOTCH4 signaling across all comparison groups, with consistently positive directionality (AGP vs. AC: z-score=3.317; VGP vs. VC: z-score=2.673; VGP vs. VGO: z-score=4.472).

This pathway enrichment was accompanied by enhanced eNOS expression in both the luminal endothelium and adventitial neovessels (Figure 6). The concurrent activation of VEGFR2-ERK-eNOS and NOTCH4 pathways—mirroring the signaling architecture that drives arterial specification during coronary embryogenesis—strongly supports the hypothesis that successful graft adaptation involves partial recapitulation of developmental arterialization programs. This coordinated pathway reactivation may represent an evolutionarily conserved adaptive mechanism whereby adult vessels redeploy embryonic programs to achieve structural and functional optimization under hemodynamic stress.

### Graft Failure is Characterized by Angiotensin-Driven Fibroinflammatory Remodeling and Loss of Biosynthetic Resilience

Our upstream regulator and regulator effects analysis identified angiotensin as a key activated node in occluded vein grafts. The angiotensin-centered network predicted a coordinated upregulation of extracellular matrix proteins (COL5A1, COL5A2, DCN, BGN, LUM, THBS2), consistent with fibrotic remodeling and reduced vascular compliance.

Immunohistochemistry confirmed increased angiotensin expression in the medial and adventitial regions of occluded vein grafts, supporting the hypothesis of an angiotensin-driven fibroinflammatory remodeling mechanism underlying graft occlusion in our cohort.

This observation aligns with mechanistic insights from the renin-angiotensin system literature: angiotensin II mediates vascular remodeling through vascular smooth muscle cell proliferation, migration, extracellular matrix production, inflammation, and oxidative stress, and is implicated in the pathogenesis of intimal hyperplasia and graft degeneration^24^.

Moreover, polymorphism studies of the angiotensin-converting enzyme (ACE) gene in saphenous vein graft atherosclerosis further support a link between angiotensin signaling and graft pathology^25^. Given that we observed significantly greater luminal deposition of angiotensin in occluded grafts compared to patent grafts, our data strengthen the hypothesis that angiotensin signaling is not merely a bystander but may function as an upstream driver of graft failure.

From a therapeutic perspective, although no large randomized trials have specifically evaluated angiotensin blockade for improving saphenous vein graft patency, preclinical and indirect clinical evidence suggest that targeting the renin-angiotensin axis may be beneficial. For example, ACE inhibitors and angiotensin receptor blockers (ARBs) reduce vascular remodeling and intimal hyperplasia in experimental models^24^. Observational analyses in post-CABG populations have examined ACE inhibitor use, with one cohort study reporting that post-CABG ACE/ARB therapy was associated with a lower hazard of major adverse cardiovascular events at 1 and 5 years^26^. Although this does not directly measure graft-specific patency, it is biologically plausible that the beneficial effect partly reflects improved graft adaptation.

Thus, in light of our finding that angiotensin accumulation is specifically associated with occluded grafts—and that this pattern corresponds to a predicted upstream regulatory network driving extracellular matrix remodeling—we propose that angiotensin signaling represents a novel, modifiable pathway in graft failure.

### Potential Role of eNOS in Perivascular Adipose Tissue as a Key Vasoprotective Modulator

eNOS is a key regulator of vascular homeostasis, mediating anti-thrombotic, anti-inflammatory, and vasodilatory effects that maintain conduit integrity. Previous clinical and experimental studies have demonstrated that increased eNOS activity within the luminal endothelium correlates with improved graft patency and reduced vascular complications in saphenous vein grafts used for coronary bypass surgery^27,28^. However, these investigations have primarily focused on endothelial eNOS localized to the main vessel lumen, without exploring its potential roles in the perivascular compartment.

In our study, we identified a distinct spatial pattern of eNOS upregulation extending beyond the endothelium to the adventitia and PVAT in patent saphenous vein grafts. Notably, intense eNOS expression was observed in the vasa vasorum and surrounding supportive structures—an expression pattern absent in pre-implantation segments. This finding suggests that the PVAT-adventitial domain may actively participate in graft adaptation, serving as a secondary vasoprotective interface that complements luminal endothelial function.

PVAT is a metabolically active layer that release paracrine mediators—such as adipokines and hydrogen sulfide—which enhance eNOS activity and exert anti-inflammatory and anti-fibrotic effects on the vascular wall. Our histological evidence supports the concept that preserving PVAT during no-touch vein harvesting not only maintains endothelial integrity but also preserves an eNOS-enriched microenvironment within the perivascular compartment. This dual-source nitric oxide signaling likely contributes to the superior long-term patency observed with the no-touch technique.

### Limitations and Future Study

This study has several limitations that warrant acknowledgment. First, the number of available graft specimens was limited, reflecting the rarity and clinical challenges of obtaining well-preserved human graft tissues explanted during cardiac transplantation. Despite the small cohort size, each sample provided valuable biological insights into long-term graft adaptation—a phenomenon rarely accessible in human studies. Second, surgical techniques were not fully standardized across cases; variations in harvesting methods and anastomotic configurations, including Y-graft constructs, may have influenced hemodynamic and remodeling responses. Third, all molecular analyses were performed on bulk tissue, precluding cell type- and layer-specific resolution. Although immunohistochemistry provided partial spatial context for key markers such as eNOS, it could not fully delineate molecular signals originating from distinct vascular compartments, including the intima, media, and adventitia.

To address these limitations, future studies should incorporate spatially resolved molecular profiling techniques—such as spatial transcriptomics and proteomics—to characterize cell type- and layer-specific remodeling signatures within arterial and venous grafts. Particular emphasis should be placed on the intimal compartment, where hemodynamic and metabolic cues converge to influence the trajectory of intimal remodeling. Spatial omics approaches will be essential for identifying the molecular regulators that determine whether intimal proliferation stabilizes adaptively or progresses toward pathological occlusion. Integrating these spatial insights with functional validation in experimental graft models will enhance mechanistic understanding and guide the development of targeted therapeutic strategies to improve long-term graft patency.

## Conclusion

In this study, we found that arterial and venous grafts with long-term patency undergo a convergent remodeling process and establish a distinct, graft-specific adaptive phenotype characterized by structural maturation, reconstitution of elastic architecture, and a shared biosynthetic-proteomic program uniquely shaped by the extraordinary hemodynamic environment of the coronary circulation. Adventitial eNOS-positive neovascularization emerged as a hallmark of adaptive remodeling, suggesting a vasoprotective mechanism that sustains graft homeostasis. Collectively, these findings identify a biosynthetic-vasoprotective remodeling axis as a central determinant of graft durability and propose that therapeutic modulation of shared adaptive pathways may enhance long-term graft patency following coronary artery bypass grafting.

## NON-STANDARD ABBREVIATIONS AND ACRONYMS

AC: Artery control
AGP: Artery graft, patent
CABG: Coronary artery bypass grafting
DAB 3: 3′-Diaminobenzidine
ECM: Extracellular matrix
eNOS: Endothelial nitric oxide synthase
FFPE: Formalin-fixed, paraffin-embedded
IGF1: Insulin-like growth factor 1
IPA: Ingenuity Pathway Analysis
LC–MS/MS: Liquid chromatography–tandem mass spectrometry
PCA: Principal component analysis
PGC-1α: Peroxisome proliferator-activated receptor gamma coactivator-1 alpha (PPARGC1A)
PPI: Protein–protein interaction
PVAT: Perivascular adipose tissue
RUNX2: Runt-related transcription factor 2
SRC: Src proto-oncogene, non-receptor tyrosine kinase
SRP: Signal recognition particle
STAT: Signal transducer and activator of transcription
VC: Vein control
VEGF: Vascular endothelial growth factor
VGO: Vein graft, occluded
VGP: Vein graft, patent

## ONLINE SUPPLEMENT

### SUPPLEMENTAL METHODS

#### Proteomic Analysis of FFPE Graft Tissues

##### Sample Preparation and Protein Extraction

For proteomic profiling, FFPE tissue sections (5 μm thick, five serial sections per sample) were prepared from 5 VGP, 3 VGO, 4 AGP, 8 VC, and 6 AC blocks and collected into LoBind tubes (Sigma-Aldrich). Samples were deparaffinized and lysed in a buffer containing 5% sodium dodecyl sulfate and 50 mM triethylammonium bicarbonate (TEAB, pH 8.5), then incubated at 50°C for 20 minutes. Sample disruption and protein extraction were performed using the Covaris R220 Focused-Ultrasonicator (Covaris, Woburn, MA). Adaptive Focused Acoustics conditions were set to 175 W, 10% duty factor, 200 cycles/burst at 20 °C for 300 seconds, followed by incubation at 80 °C for 1 h.

Reduction (20 mM dithiothreitol, 56°C for 10 minutes) and alkylation (40 mM iodoacetamide, 30 minutes in the dark at room temperature) were performed, followed by a second acoustic treatment (360 seconds). Protein concentrations were measured via Bradford assay (Thermo Fisher). Protein digestion was carried out using the Suspension Trap (S-Trap, Profiti) protocol. After acidification with phosphoric acid and binding in methanol/TEAB buffer, proteins were trapped, washed, and sequentially eluted with TEAB, 0.2% formic acid, and 0.2% formic acid/50% acetonitrile. The eluates were vacuum-dried and stored at –80°C.

##### LC-MS/MS Acquisition and Protein Identification

Peptides were analyzed using a Vanquish Neo LC system coupled with an Orbitrap Eclipse mass spectrometer (Thermo Fisher Scientific). Samples (4 μL, 0.5 μg/μL) were injected into a PepMap C18 trap column and separated on a PepMap Neo analytical column (75 μm × 500 mm, 2 μm, 100 Å) at a flow rate of 300 nL/min over 180 minutes. A gradient of 3–25% solvent B over 150 minutes, 25–80% B over 10 minutes, and 80% B for 20 minutes was applied (mobile phases: 0.1% formic acid in water or acetonitrile). The Orbitrap operated in data-dependent acquisition mode with MS1 resolution set to 120,000, scan range of 375–1800 m/z, and FAIMS compensation voltages of –45, –55, and –75 V. MS2 spectra were acquired in the ion trap using turbo scan mode (AGC target 300%, maximum injection time 15 ms, normalized collision energy 35%).

##### Database Search and Quantification

Raw files were analyzed using SEQUEST-HT in Proteome Discoverer v2.4 against the UniProt-SwissProt human database. Search parameters included a 10 ppm precursor tolerance and a 0.6 Da fragment tolerance, allowing for two missed cleavages. Carbamidomethylation was set as a fixed modification, and methionine oxidation as a variable modification. The false discovery rate (FDR) was estimated using a target-decoy strategy and controlled at <1% using Percolator.

Label-free quantification (LFQ) was performed based on precursor ion intensities of unique and razor peptides, excluding oxidized methionine. LFQ values were log2-transformed, normalized using width adjustment, and imputed with a normal distribution model. Proteins detected in all replicates of at least one group were retained. Principal component analysis (PCA) was conducted using Perseus (v1.6.2.3).

Data visualization and exploratory clustering analyses were performed with Instant Clue, including volcano plots, correlation plots, hierarchical heatmaps, and cluster box plots. Hierarchical clustering was carried out using Distance=Correlation and Linkage=Complete.

Sparse partial least squares–discriminant analysis (sPLS-DA) was performed using MetaboAnalyst (www.metaboanalyst.ca). Upset plots were generated with the ComplexUpset R package (v1.3.3) implemented in RStudio (2024.12). Pathway enrichment analyses were conducted using Metascape (https://metascape.org) and Ingenuity Pathway Analysis (IPA, QIAGEN Inc.).

##### Canonical Pathway Analysis

Canonical pathway enrichment was conducted using the Core Analysis function in IPA (QIAGEN) on differentially expressed proteins (DEPs) between graft groups. The directional interpretation of pathway activity was based on the IPA z-score metric, which integrates observed protein expression changes with curated activation and inhibition relationships from the IPA knowledge base. A positive z-score indicates a pathway predicted to be activated, while a negative z-score suggests inhibition. Pathways with |z| ≥ 2 were considered to have strong predictive directionality. Pathways lacking sufficient data for z-score calculation were excluded from visual summaries. Visual heatmaps represent these z-scores using color scales centered at zero, with blue indicating suppression and orange indicating activation.

##### Immunohistochemistry and Quantitative Image Analysis

Immunohistochemical staining was performed using specific antibodies against angiotensinogen, the precursor of angiotensin, C-Myc, EGFR, IGF1/IGF2, NKX2.3, NR4A3 (NOR-1), PGC-1α (PPARGC1A), PTGES, RETN (Resistin), RUNX2, SRC, STAT1, and phospho-STAT3 (Tyr705) to evaluate protein expression in graft tissues. All antibodies were validated for FFPE human tissues, and staining was conducted according to the manufacturers’ protocols, with minor optimizations for antigen retrieval and detection conditions.

##### Endothelial and Vascular Marker Staining

Immunohistochemistry was performed using antibodies against CD31, α-smooth muscle actin (αSMA), and endothelial nitric oxide synthase. All stained slides were digitized using a whole-slide scanner (Aperio GT450 DX, Leica Biosystems), and image analysis was conducted using QuPath (v0.5.1). Detailed information on antibody sources, catalog numbers, clones, and working dilutions is provided in Table S2.

##### eNOS Signal Quantification in Endothelial Cells

Using the area annotation tool in QuPath, the innermost luminal layer was manually annotated to select the target area for analysis. Nuclear segmentation was then performed based on the hematoxylin signal. Because stromal cells near the lumen were occasionally included in this segmentation, manual curation was conducted to remove any non-endothelial cells, ensuring that only true endothelial cells were segmented. Specifically, cells exhibiting characteristic hobnail morphology, indicative of true luminal endothelial cells, were retained, while cells located deeper within the stroma were excluded, as shown in Figure S1C-D.

For each nucleus, the mean optical density of the DAB signal for eNOS was measured.

A histogram of per-cell eNOS intensity was generated to visualize the distribution. A threshold of 0.2 was applied to distinguish eNOS-positive from eNOS-negative nuclei. The proportion of eNOS-positive endothelial cells was then calculated relative to the total number of segmented endothelial cells.

##### Quantification of CD31-Positive Neovessels

To assess neovascularization, including vasa vasorum extending from the media to the adventitia, CD31-stained sections were analyzed using a segmentation-based approach in QuPath. The parameters included a spatial resolution of 0.51 μm/pixel, a Gaussian filter value of 2, an intensity threshold of 0.44, and object/hole size thresholds of 20 pixels. This pipeline enabled consistent detection of small capillaries (Figure S1E–H). Quantification metrics included the total capillary area and the cumulative perimeter of CD31-positive structures.

##### Digital Quantification of IHC Using QuPath

Immunohistochemical signals were quantified on whole-slide images using QuPath (v0.5.1). For each slide, regions of interest (ROIs) were manually annotated to include representative lesional areas while excluding artifacts. Stain vectors (hematoxylin and DAB) were calibrated on the same slide prior to analysis. Nuclei were segmented using the “Positive cell detection” workflow, with nuclear DAB optical density (OD) serving as the scoring compartment. Intensity thresholds were established to classify cells into 0, 1+, 2+, and 3+ categories consistently across slides for each antibody. Measurements were exported at the annotation level.

For each ROI, the positive percentage was calculated as (#1+ + #2+ + #3+) divided by the total number of nuclei, multiplied by 100. The H-score[1] was computed on a scale ranging from 0 to 300. All thresholding and post-processing parameters were kept constant within each marker to ensure comparability across study groups.

##### Statistical Analysis

All statistical analyses were conducted using R (v3.2.3; R Foundation for Statistical Computing). Group comparisons were performed using analysis of variance (ANOVA). To account for repeated measures and inter-individual variability, generalized estimating equations (GEE) were applied, implemented via the geeglm function in the geepack package. Post hoc pairwise comparisons were adjusted using Tukey’s method, with Bonferroni correction applied to control for multiple testing.

Raw data from QuPath analysis were imported and processed using Python 3.8 with pandas (version 1.5.3) for data manipulation. A total of 12 proteins were analyzed: angiotensinogen, c-MYC, EGFR, IGF1, NR4A3, PTGES, PGC, RETN, RUNX2, SRC, STAT1, and STAT3.

Quantitative immunohistochemical data were analyzed using non-parametric statistical methods due to the data’s non-normal distribution. For each protein, two primary outcome measures were evaluated: the percentage of positive cells (Positive %) and the H-score, both derived from QuPath analysis software. Statistical significance was set at α=0.05.

Results were visualized using box-and-whisker plots combined with individual data point overlays (swarm plots) to illustrate both distribution characteristics and individual observations. The plots were generated using the Matplotlib (version 3.7.1) and Seaborn (version 0.12.2) libraries, adhering to academic formatting standards suitable for publication. Statistical significance was indicated on the plots with horizontal lines and asterisk notation above the compared groups.

## SUPPLEMENTAL RESULTS

### Histomorphological Characteristics of Patent Arterial and Venous Grafts (continued)

Native arterial conduits, freshly harvested for CABG (AC), exhibited the expected features of elastic arteries, including a compact medial layer rich in elastic fibers and well-defined internal and external elastic laminae. After grafting, AGP retained this elastic-artery framework but showed reinforcement of the internal elastic lamina and marked thickening of the muscular media, with minimal intimal hyperplasia—findings consistent with stable structural adaptation. Native venous conduits, freshly harvested for CABG (VC), exhibited thin vessel walls, wide lumina, sparse medial elastic fibers, and an indistinct elastic laminar structure.

Quantitative morphometric analysis revealed distinct graft-type–specific patterns in vessel dimensions (Figure 1C). Lumen area showed minimal differences between AGP and their native AC. VGP exhibited a modest reduction in lumen area compared with VC, resulting in lumen dimensions that approached those of AGP—reflecting the extent to which the venous conduit had remodeled toward the size range of an arterial conduit. In contrast, VGO showed a pronounced increase in lumen area (P<0.001), consistent with the aneurysmal dilatation observed histologically. A similar pattern was observed in lumen circumference, with VGO demonstrating substantial enlargement relative to all other groups.

Intimal measurements revealed mild thickening in AGP compared with AC (P<0.001), indicating neointimal growth in arterial grafts. VGO showed a markedly enlarged intimal area compared with VGP (P<0.001), consistent with the severe structural distortion observed in failed grafts. Medial thickness was significantly greater in AGP than in AC (P<0.001), reflecting adaptive medial hypertrophy. VGP showed a non-significant trend toward increased medial thickness relative to VC. Conversely, VGO exhibited a significant reduction in medial thickness compared with both VGP and VC (P<0.001), indicating medial degeneration associated with graft failure.

### Intimal and Medial Morphology Across Arterial and Venous Graft States (Figure 1D)

In AGP, intimal changes were minimal compared with native AC. AGP showed a well-preserved luminal surface without cleft formation, a more prominent internal elastic lamina, and reduced elastic fiber content within the media—features indicative of a shift from an elastic artery pattern toward a more muscular artery configuration.

VGP exhibited substantial intimal thickening compared with native VC. The intima of VGP contained interspersed clefts and internal striations, accompanied by a markedly thickened internal elastic lamina. The media showed increased elastic fiber deposition along with smooth muscle hypertrophy, indicating progressive structural remodeling in response to arterial hemodynamic stress. In contrast, VGO exhibited a distinctly different pattern. The intimal surface was smooth and rounded, lacking clefts, and was composed predominantly of dense, disorganized collagen. The endothelial lining was absent, and luminal calcification was frequently observed. The media showed extensive necrosis and loss of normal elastic laminar structures. These findings highlight that, although both VGP and VGO develop intimal thickening, the underlying organization and associated medial changes differ profoundly between patent and failed grafts.

### Robust Proteomic Profiling of Archival Graft Tissue

To evaluate protein expression signatures across graft types, we performed label-free quantitative proteomics on FFPE vascular grafts. Across all 26 samples, an average of over 3,000 proteins were confidently identified per sample (Figure S2). Quantile-based normalization (Q1–Q3) of protein abundance resulted in tightly aligned log₂ distributions, minimizing systematic bias and enhancing cross-sample comparability. Following normalization, protein abundance profiles exhibited nearly identical ranges and medians across groups. Technical reproducibility was confirmed by Pearson correlation analysis, which revealed strong within-group clustering and high concordance between biological replicates (Figure S2).

A total of 4,836 proteins were identified across all vascular samples. Among these, 4,012 proteins were shared across all five groups, representing a substantial common vascular proteome (Figure S3).

### Sparse Partial Least Squares Discriminant Analysis

To show group separation, we performed sPLS-DA (Figure S3B). Consistent with the PCA findings, AGP and VGP showed close proximity, whereas VGO remained distinctly segregated from all other groups. To identify discriminative molecular features, we examined the loading profiles. The highest-ranked contributors on Component 1 included glycolytic enzymes, cytoskeletal regulators, and stress-associated proteins.

The three-dimensional plot showed that both AGP and VGP clustered distinctly from their respective controls, indicating convergent remodeling signatures under arterial hemodynamic stress. In contrast, VGO samples formed a separate cluster, distinct from both VGP and VC, reflecting a proteomic profile consistent with maladaptive or fibro-inflammatory remodeling.

Examination of the loading vectors identified the proteins most strongly contributing to Component 1. High VIP scores were observed for proteins that regulate glycolysis, cytoskeletal stability, and mitochondrial protein homeostasis, which were consistently elevated in functional grafts (AGP and VGP), reflecting active metabolic remodeling and preservation of structural integrity under arterial hemodynamic stress.

Component 2 captured vein-specific remodeling features driven by extracellular matrix and mitochondrial regulatory proteins. These proteins were enriched in VGO relative to AGP/VGP, indicating enhanced ECM deposition, altered mitochondrial signaling, and immune-activated remodeling characteristic of graft failure.

### Divergent Baseline Proteomic Programs in Native Arteries and Veins (Clusters 6 and 7)

Unsupervised hierarchical clustering identified two native vessel–specific clusters, corresponding to Cluster 6 (AC) and Cluster 7 (VC). Pathway enrichment analysis (Figure S4A) revealed that Cluster 6 (AC) was enriched in biological processes related to epithelial differentiation, keratinization, and intermediate filament organization. In contrast, Cluster 7 (VC) showed preferential enrichment of pathways associated with vascular smooth muscle contraction, actin cytoskeleton remodeling, and extracellular matrix interactions. PPI network analysis (Figure S4B) further supported these distinctions: AC networks were dominated by keratin family members, laminins, and collagens, whereas VC networks centered on actomyosin contractile proteins. Metascape enrichment bubble plots (Figure S4C) highlighted these contrasting themes, with AC associated with structural reinforcement and barrier-related programs, and VC linked to cytoskeletal plasticity, cell–matrix adhesion, and vascular repair pathways. Together, these findings reinforce the baseline molecular divergence between native arteries and veins and provide a reference framework for interpreting graft-associated remodeling.

### Upstream Regulators and Pathway Programs Distinguishing Arterial Grafts from AC

Upstream regulator analysis comparing AGP with native internal AC revealed a coordinated upregulation of multiple metabolic and proliferative regulators (Figure 4A). Key activated upstream nodes included IGF1, RUNX2, NR4A3, PGC-1α, EGFR, and MYC, which converge on biological functions related to cell proliferation, connective tissue growth, and metabolic activation. These regulators formed a coherent signaling network, indicating that arterial grafts engage a distinct adaptive remodeling program absent in control arteries.

Consistent with these upstream predictions, AGP demonstrated enrichment of canonical pathways supporting the biosynthetic and metabolic aspects of vascular remodeling (Figures S6–S10). These pathways included those mediating protein quality control and stress adaptation (BAG2 signaling), mitochondrial energy metabolism (oxidative phosphorylation, sirtuin signaling), biosynthetic translational control (mTOR signaling, regulation of eIF4 and p70S6K), autophagy and vesicular turnover (microautophagy signaling), innate immune effector pathways (neutrophil degranulation, nitric oxide and reactive oxygen species production), angiogenic signaling (VEGF pathway), and nutrient stress–responsive translational regulation (signal recognition particle [SRP]–dependent cotranslational targeting, EIF2AK4/GCN2 amino acid deficiency response, EIF2 signaling).

Collectively, these findings show that arterial graft adaptation is characterized by IGF1–RUNX2–driven metabolic and proliferative reprogramming, supported by coordinated activation of mitochondrial, biosynthetic, and immunoregulatory pathways. This integrated signaling architecture defines an adaptive remodeling phenotype distinct from the quiescent state of native arterial conduits.

### Activation of Proliferative and Biosynthetic Programs and Suppression of Contractile

#### Regulators in Patent Vein Grafts (continued)

Pathway-level analyses (Figures S11–S13) further substantiated the findings shown in Figure 4B. VGP demonstrated reduced activity in the GP6 signaling pathway, a collagen receptor cascade, suggesting decreased platelet–extracellular matrix interactions during vein graft adaptation. Concurrent alterations were observed in components of collagen biosynthesis and modifying enzyme networks, reflecting extracellular matrix remodeling associated with vein graft adaptation to arterial hemodynamics. VGP also exhibited enrichment of EIF2 signaling and integrin signaling pathways, indicating activation of translational control and cytoskeletal remodeling programs. Additionally, VGP showed activation of signal recognition particle (SRP)–dependent cotranslational protein targeting, the EIF2AK4/GCN2 amino acid deficiency response, and the integrated stress response. These translational stress-adaptation pathways overlapped with those activated in arterial grafts (AGP vs. AC), suggesting conserved protein synthesis and stress-response programs across both graft types during adaptation to arterial pressure environments.

## SUPPLEMENTAL FIGURES

**Figure S1.**
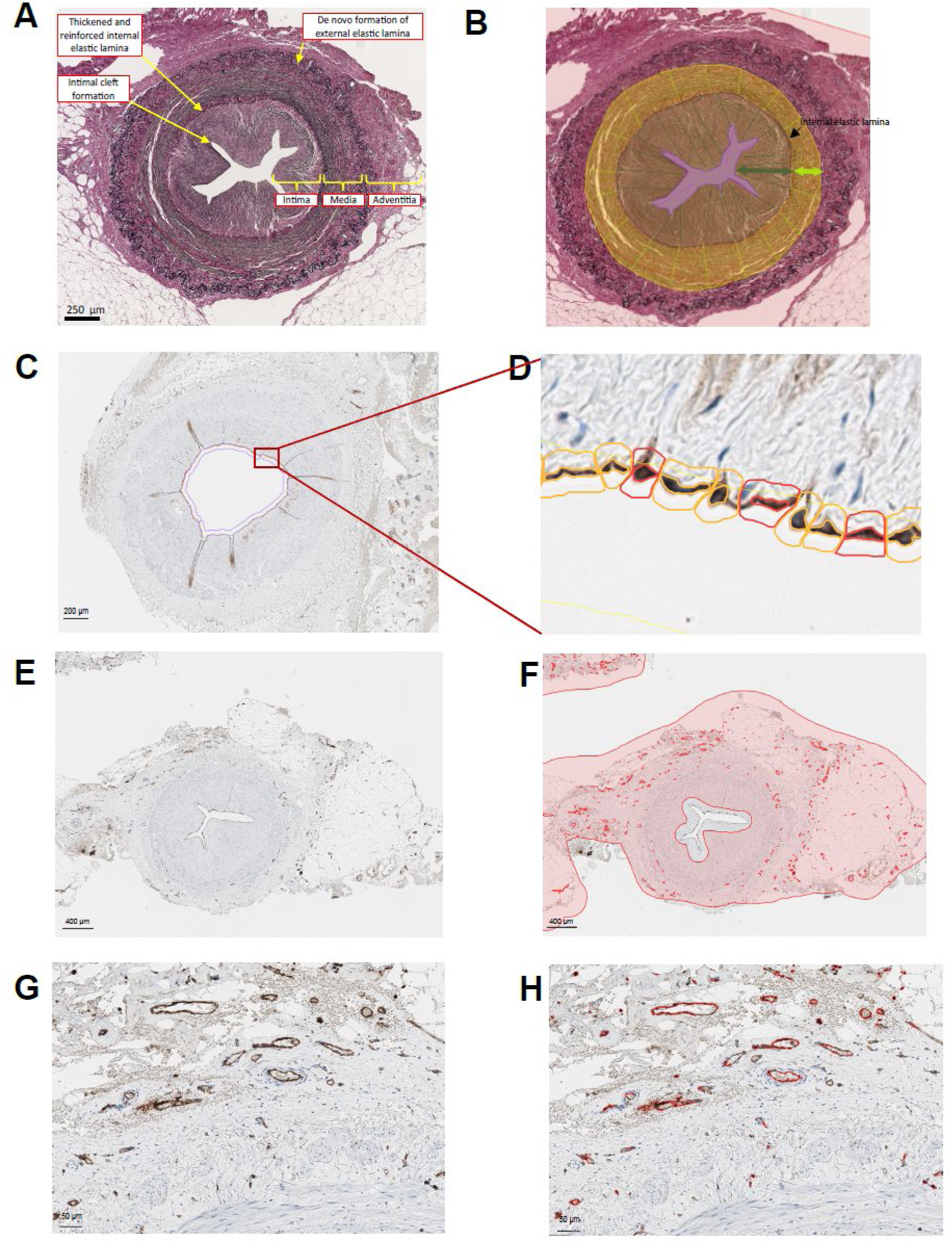
(A) Long-term patent SV graft showing significant intimal hyperplasia and cleft formation Histomorphological analysis of long-term patent SV grafts revealed intimal hyperplasia, cleft formation, and increased eNOS expression in the luminal endothelium and adventitial vasa vasorum. (B) Digitally scanned images were analyzed using QuPath with manual annotation and quantification. The lumen area (L, purple line) was measured by delineating the innermost lumen perimeter. Intimal thickness (I, green arrow) was defined as the distance between the internal elastic lamina (IEL, black line) and the main lumen, while medial thickness (M, light green arrow) was measured as the distance between the IEL and the outer border of the medial smooth muscle layer. Cleft depth (magenta line) was defined as the distance from a virtual line smoothing the main lumen to the deepest part of the cleft. (C-D) Digital image analysis workflow for quantifying luminal eNOS. C, Representative eNOS immunohistochemistry with annotated luminal endothelium (pink outline). Scale bar=200 µm. D, High-magnification view demonstrating QuPath-based nuclear segmentation (hematoxylin, blue) and cell boundary detection (orange/red outlines). Mean DAB optical density was quantified for each nucleus across 20,792 endothelial cells for single-cell distribution analysis (Figure 6B). (E-H) Quantification of adventitial CD31-positive capillaries. Whole-section view showing CD31-positive vasa vasorum. Scale bar=400 µm. D, Annotation of the media-adventitial compartment (pink) for selective microvessel analysis, excluding the luminal endothelium. Scale bar=400 µm. (G-H) High-magnification view of CD31-positive adventitial neovascularization in a patent saphenous vein graft. Scale bar=50 µm. F, QuPath segmentation of CD31-positive vessels (red outlines) for automated perimeter quantification (Figure 6E, left). Scale bar=50 µm. All sections were digitized at 20× magnification. QuPath (v0.4.3) parameters included a nuclear diameter of 5–15 µm, cytoplasmic expansion of 2 µm, and DAB color deconvolution using standard vectors. The eNOS positivity threshold was defined as an optical density greater than 0.2.

**Figure S2.**
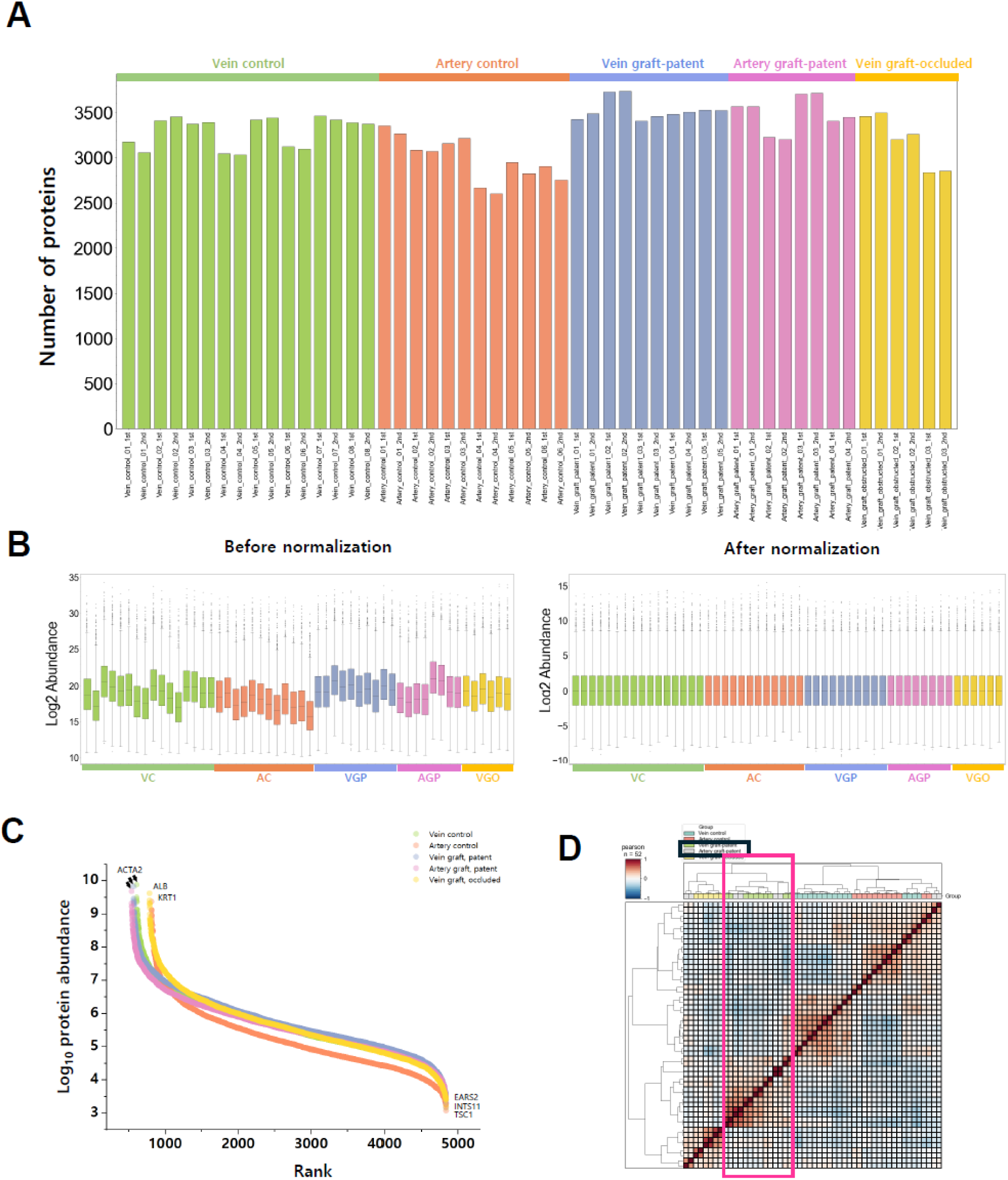
Robust proteomic profiling of archival vascular graft tissue (A) Number of identified proteins. Bar plot showing the total number of confidently identified proteins per sample across all graft categories. Despite the archival formalin-fixed, paraffin-embedded nature of the tissues, each sample yielded high-depth proteomic coverage, with an average of over 3,000 proteins identified. (B) Distribution of sample abundance. Distribution of log₂-transformed protein abundance values before (left) and after (right) quantile-based normalization (Q1–Q3). Each box plot represents individual samples grouped by vessel type: VC (native vein controls), AC (native arterial controls), VGP (patent venous grafts), AGP (patent arterial grafts), and VGO (occluded venous grafts). Before normalization, substantial systematic variability is evident across groups, characterized by differing medians and dynamic ranges. After normalization, the distributions were aligned, exhibiting comparable medians and interquartile ranges across all groups; this process minimizes technical bias while preserving biological differences. The normalization method employed was asymmetric quartile-based scaling, in which the median (Q2) was subtracted from all values to center each distribution; values above the median were scaled by (Q3 − Q2), and values below the median were scaled by (Q2 − Q1). (C) Ranked protein abundance. Rank–abundance plots demonstrating consistent detection of highly abundant vascular structural proteins, including ACTA2, KRT1, and albumin, across all samples and graft types. (D) Pearson correlation heatmap revealed strong within-group clustering and high concordance between biological replicates, confirming the robust technical reproducibility of the proteomic dataset.

**Figure S3.**
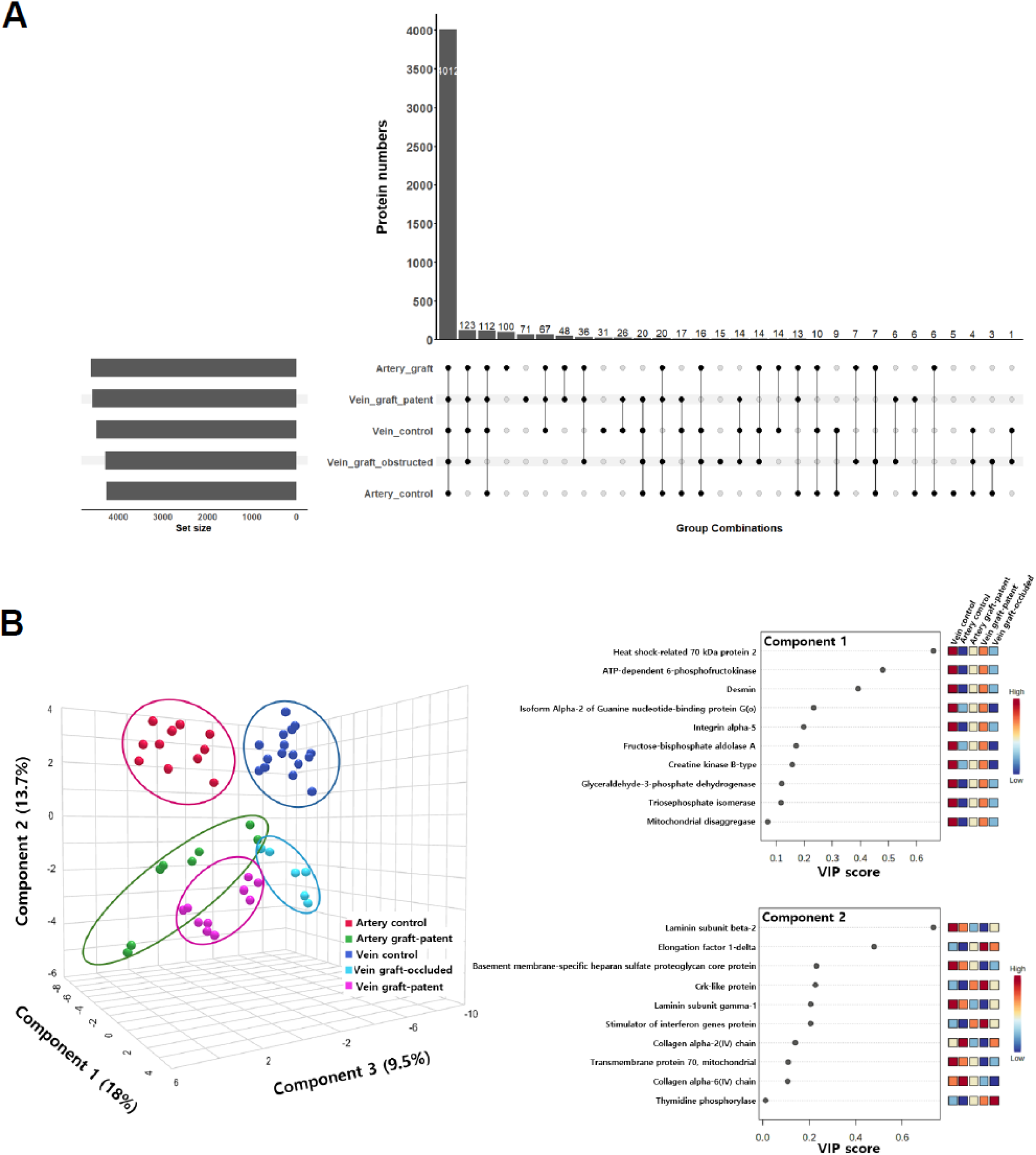
(A) Intersection analysis of detected proteins across vascular groups using the ComplexUpset package (v1.3.3). A total of 4,836 proteins were identified across all samples. The bar plot on the left displays the number of detected proteins within each group (AC, VC, AGP, VGP, VGO), demonstrating comparable proteomic depth across the datasets. The intersection matrix and the accompanying bar plot illustrate the shared and unique protein sets among the five groups, with 4,012 proteins commonly detected across all groups. Each group also exhibits a subset of uniquely detected proteins, indicating condition-specific proteomic features that distinguish native vessels, patent grafts, and occluded grafts. (B) sPLS-DA clustering and discriminant protein loadings. (Left) 3D sPLS-DA score plot (Components 1–3) illustrating the separation among native vessels (AC, VC), patent grafts (AGP, VGP), and occluded grafts (VGO). AGP and VGP showed partial overlap, whereas VGO formed a discrete cluster. (Right) Loading plots for Components 1 and 2, highlighting proteins with the greatest contribution to group discrimination. The top-ranked contributors include glycolytic enzymes (PFKP, ALDOA, GAPDH), cytoskeletal proteins (DES, ITGA5), and stress-associated markers (HSPA2, CKB).

**Figure S4.**
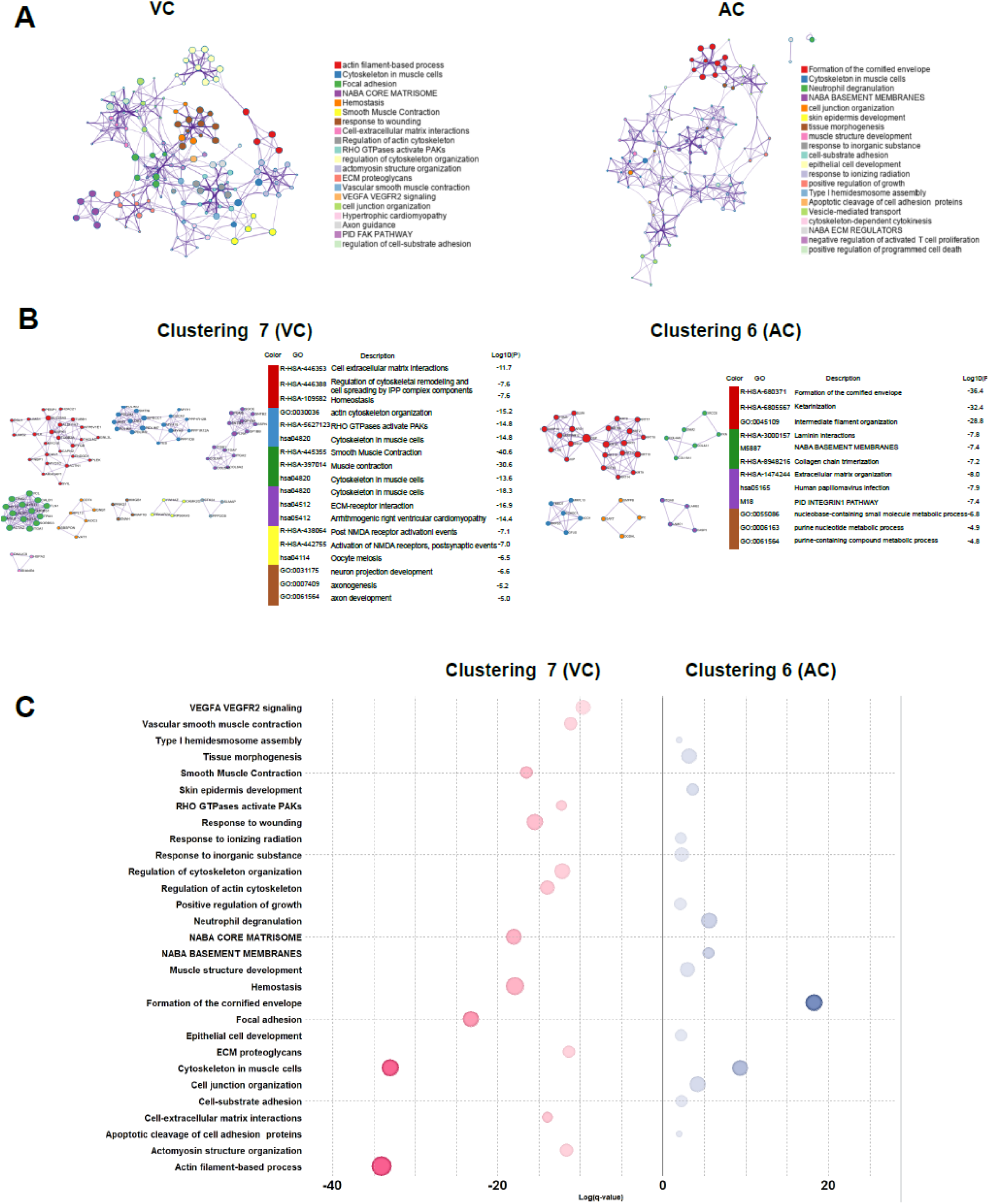
Baseline proteomic signatures of native arteries and veins (Clusters 6 and 7). (A) PPI network maps for Cluster 6 (AC) and Cluster 7 (VC), illustrating distinct interaction modules and dominant protein groups within each native vessel cluster. (B) Metascape functional enrichment networks showing major biological processes represented in Cluster 6 and Cluster 7. (C) Metascape enrichment bubble plots (GO/KEGG/Reactome) highlighting the top enriched pathways for each cluster. Cluster 6 (AC) was enriched for epithelial differentiation, keratinization, and intermediate filament organization, whereas Cluster 7 (VC) was enriched for vascular smooth muscle contraction, actin cytoskeleton remodeling, and ECM-associated processes. These analyses confirm the divergent baseline proteomic programs of arteries and veins, providing a reference framework for interpreting graft-associated remodeling.

**Figure S5.**
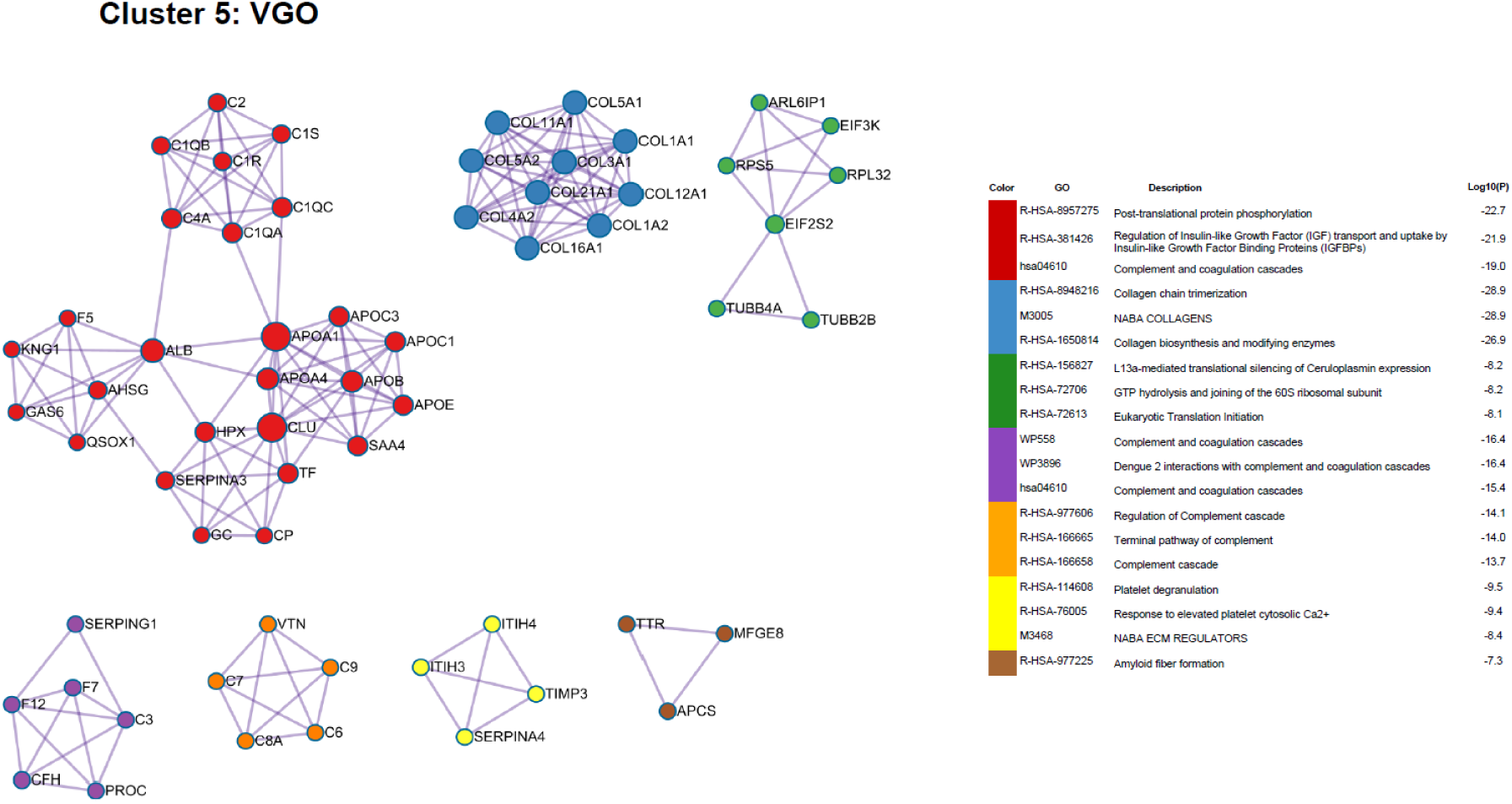
Protein–protein interaction (PPI) networks for Cluster 5. Cluster 5 PPI networks formed extensive complement and coagulation clusters, indicative of a maladaptive and pro-thrombotic remodeling program.

**Figure S6.**
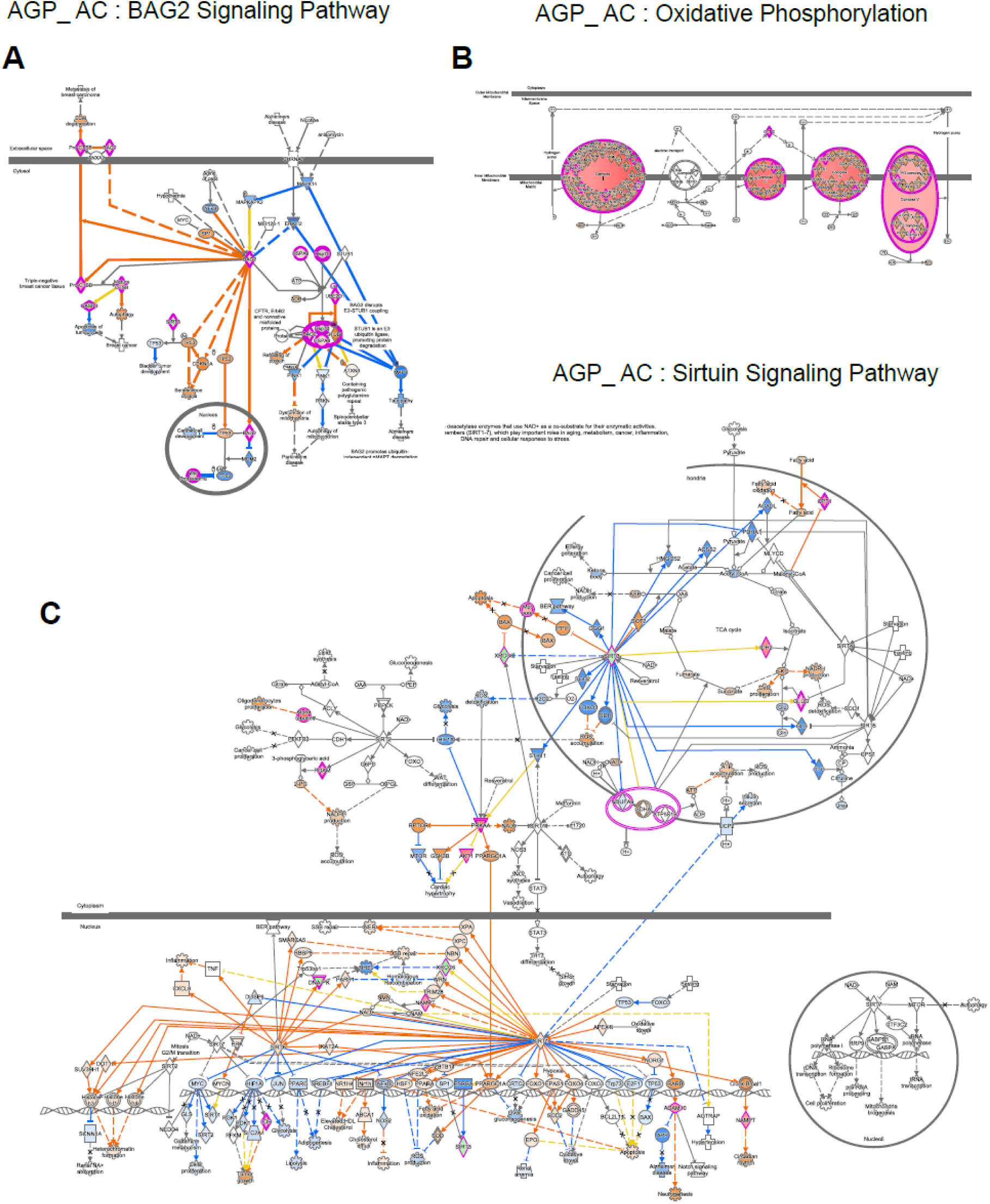
IPA canonical pathway maps highlighting BAG2 signaling, oxidative phosphorylation, and sirtuin signaling in arterial grafts (AGP vs. AC). (A) BAG2 signaling pathway. Ingenuity Pathway Analysis (IPA) canonical pathway diagram showing upregulated components (magenta) involved in BAG2-mediated chaperone regulation and proteostasis in AGP relative to AC. The predicted activation flow (orange arrows) indicates enhanced protein quality control and chaperone-dependent degradation signaling. (B) Oxidative phosphorylation. Canonical pathway map illustrating the increased representation of mitochondrial respiratory chain components in AGP. Highlighted nodes (magenta) correspond to complexes involved in electron transport and ATP production, consistent with the elevated mitochondrial oxidative capacity observed in arterial grafts. (C) Sirtuin signaling pathway. IPA pathway diagram demonstrating the differential activation of sirtuin-associated metabolic and stress-response modules in AGP. Upregulated nodes (magenta) and predicted activation steps (orange and blue arrows) reflect enhanced mitochondrial metabolism, redox homeostasis, and NAD⁺-dependent regulatory signaling in grafted arterial tissue.

**Figure S7.**
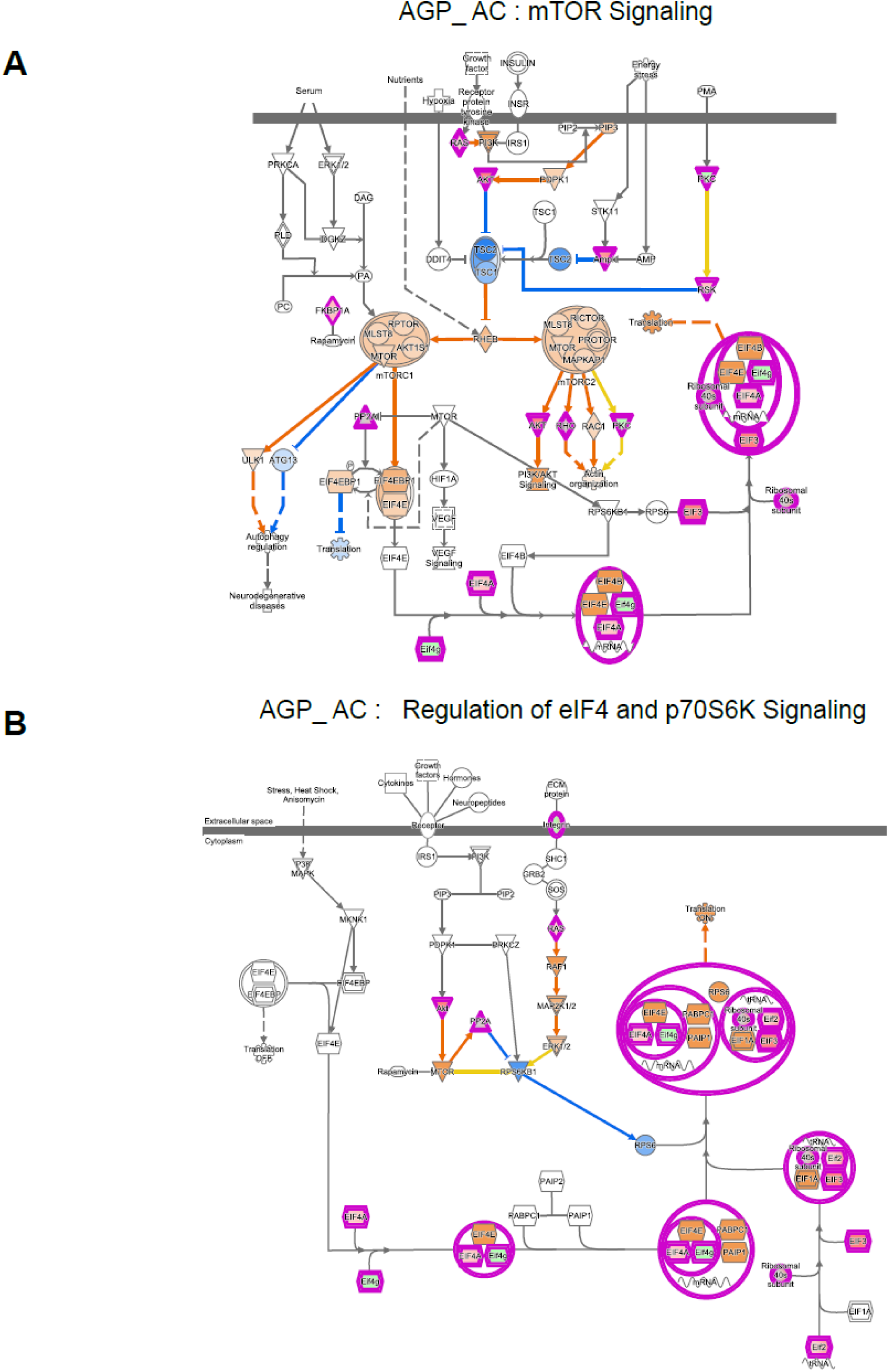
IPA canonical pathway maps highlighting mTOR signaling and translational control in arterial grafts (AGP vs. AC). (A) mTOR Signaling. Ingenuity Pathway Analysis (IPA) canonical pathway diagram showing differential activation of mTORC1- and mTORC2-associated modules in AGP relative to AC. Upregulated nodes (highlighted in magenta) and predicted activation flow (orange arrows) reflect enhanced nutrient- and growth factor–responsive signaling, including the downstream regulation of protein synthesis, autophagy, and cytoskeletal organization. (B) Regulation of eIF4 and p70S6K Signaling. Canonical pathway map illustrating the upregulation of translation initiation machinery and ribosomal protein activation in AGP versus AC. Highlighted nodes (magenta) and predicted activation steps (orange arrows) indicate increased eIF4 complex activity, p70S6K signaling, and translational output, consistent with the elevated biosynthetic demand in arterial grafts.

**Figure S8.**
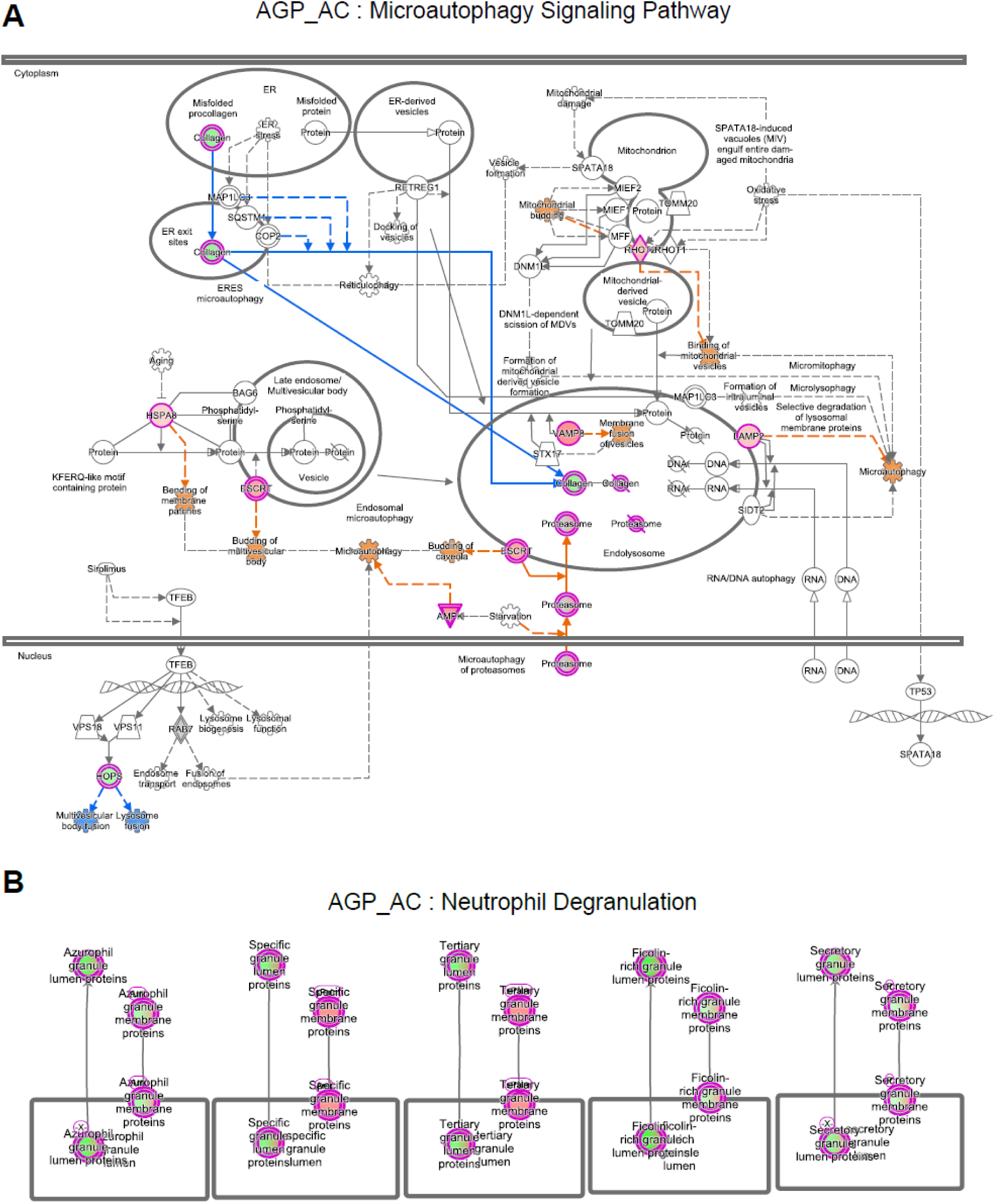
IPA canonical pathway maps illustrating microautophagy signaling and neutrophil degranulation in arterial grafts (AGP vs. AC). (A) Microautophagy Signaling Pathway. Ingenuity Pathway Analysis (IPA) canonical pathway diagram showing upregulated components (highlighted in magenta) involved in ER-derived vesicle formation, endosomal and lysosomal trafficking, mitophagy-related processes, and microautophagy-associated protein turnover in AGP relative to AC. Predicted activation flows (orange arrows) reflect enhanced autophagy-linked proteostasis mechanisms in arterial grafts. (B) Neutrophil Degranulation. Canonical pathway map depicting proteins associated with azurophilic, specific, tertiary, and secretory granule compartments. Highlighted nodes (magenta) represent granule-associated proteins upregulated in AGP compared with AC, indicating enhanced neutrophil degranulation–related signatures within the arterial graft microenvironment.

**Figure S9.**
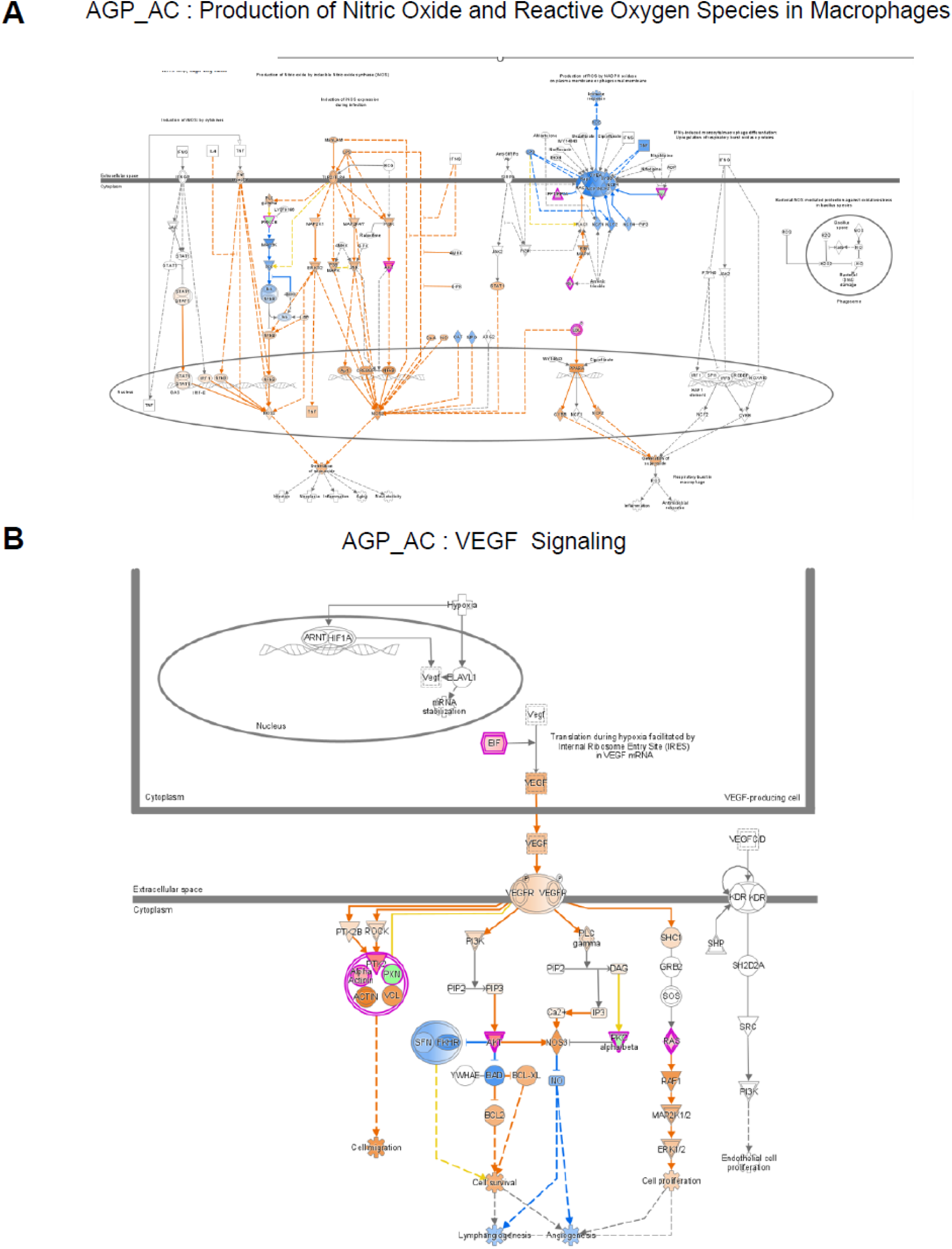
IPA canonical pathway maps highlighting nitric oxide/ROS production and VEGF signaling in arterial grafts (AGP vs. AC). (A) Production of Nitric Oxide and Reactive Oxygen Species in Macrophages. Ingenuity Pathway Analysis (IPA) canonical pathway diagram showing differential activation of nitric oxide– and reactive oxygen species–related signaling components in AGP relative to AC. Upregulated nodes (highlighted in magenta) and predicted activation flow (orange arrows) illustrate enhanced redox-associated signaling in arterial grafts. (B) VEGF Signaling. Canonical pathway map depicting VEGF signaling in the AGP vs. AC comparison. Highlighted nodes (magenta) and predicted activation pathways (orange arrows) reflect enhanced VEGF receptor–mediated signaling, including downstream PI3K/AKT, MAPK/ERK, and endothelial survival/migration pathways.

**Figure S10.**
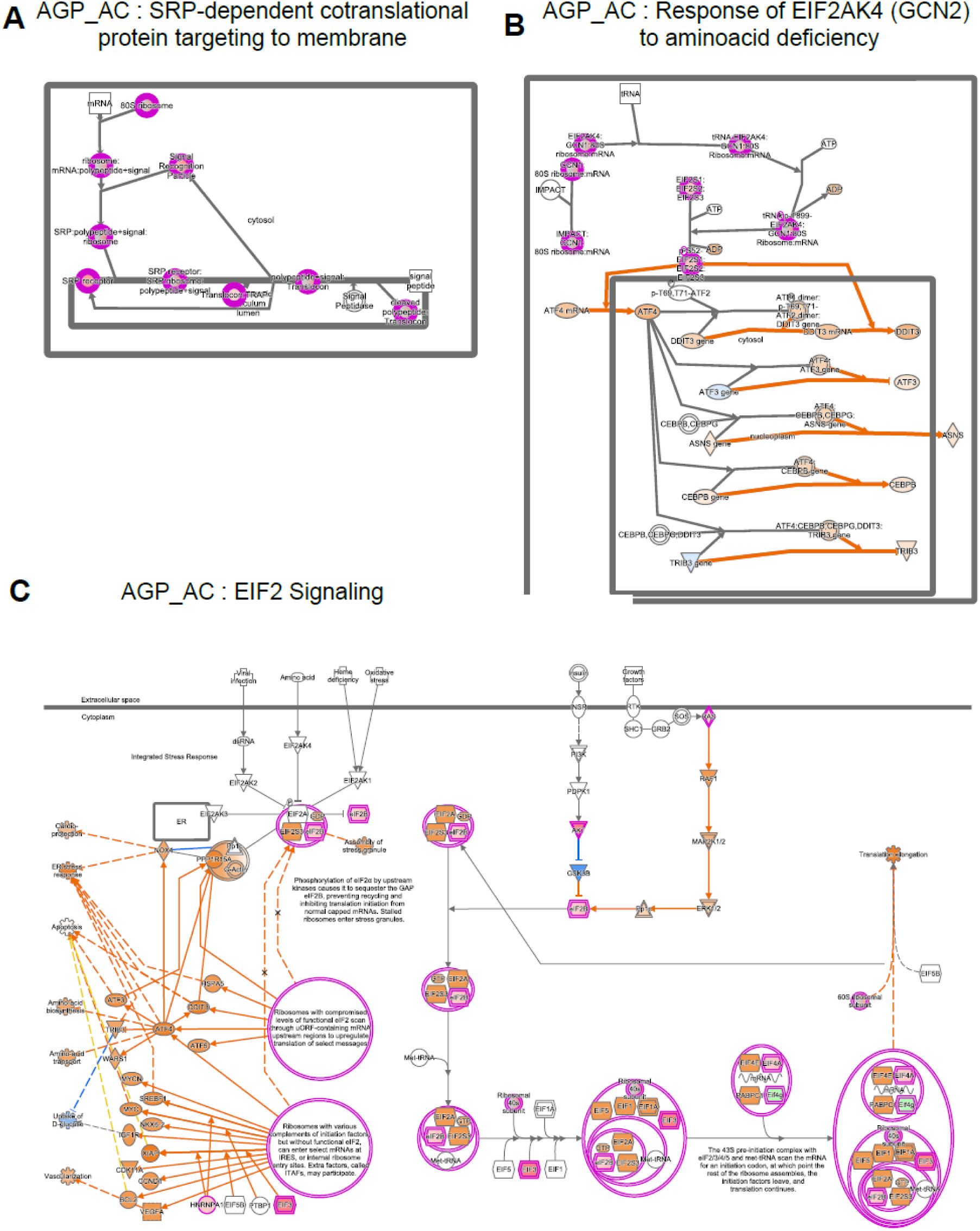
IPA canonical pathway maps highlighting translational and stress-response signaling in arterial grafts (AGP vs. AC). (A) SRP-dependent cotranslational targeting of proteins to the membrane. Ingenuity Pathway Analysis (IPA) canonical pathway diagram showing upregulated components (magenta) in AGP relative to AC, indicating enhanced engagement of ribosome–SRP complexes and ER-directed cotranslational processing. (B) Response of EIF2AK4 (GCN2) to amino acid deficiency. Canonical pathway map illustrating the activation of the GCN2–ATF4 axis in AGP. Upregulated nodes (magenta) and predicted activation flow (orange arrows) highlight enhanced amino acid sensing and downstream integrated stress response gene activation. (C) EIF2 signaling. Pathway diagram illustrating the broad activation of EIF2 signaling in AGP, including upregulated translation initiation factors, ribosomal proteins, and ATF-dependent stress-response modules. Predicted activation (orange arrows) and highlighted nodes (magenta) reflect increased translational demand and ISR engagement in arterial grafts.

**Figure S11.**
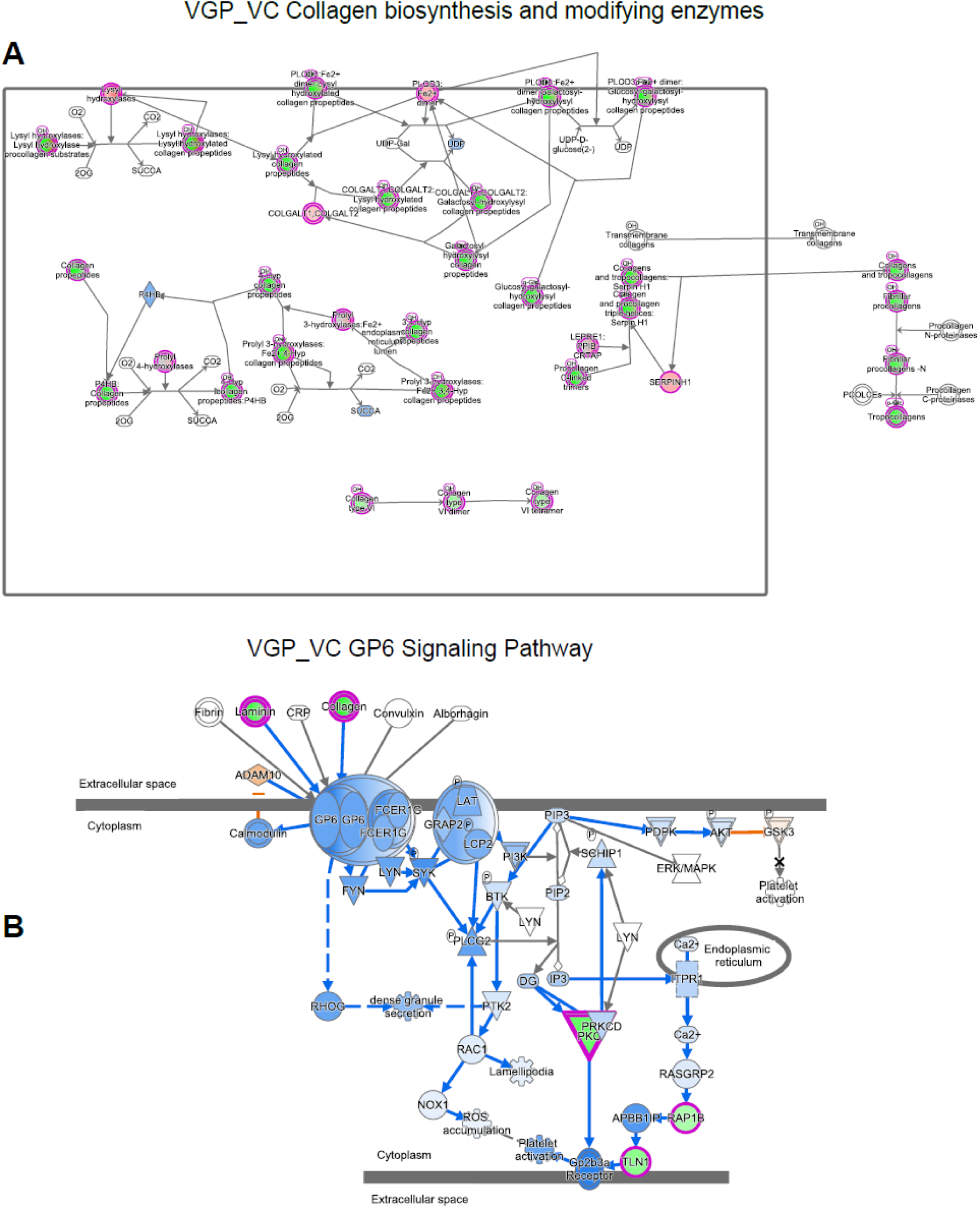
IPA canonical pathway maps illustrating extracellular matrix remodeling and platelet-related signaling in functional vein grafts (VGP vs. VC). (A) Collagen biosynthesis and modifying enzymes. Ingenuity Pathway Analysis (IPA) canonical pathway map showing upregulated components (magenta) involved in collagen chain synthesis, hydroxylation, glycosylation, cross-linking enzymes (e.g., LOX family), and procollagen processing in VGP relative to VC. These enriched nodes highlight enhanced extracellular matrix production and collagen maturation in functional vein grafts. (B) GP6 signaling pathway. IPA canonical pathway map for GP6 signaling demonstrating altered platelet-related signaling events in the VGP vs. VC comparison. Upregulated nodes (magenta) and predicted activations (blue/orange arrows depending on pathway direction) indicate changes in collagen–platelet interaction modules, downstream cytoskeletal rearrangements, and platelet activation signaling.

**Figure S12.**
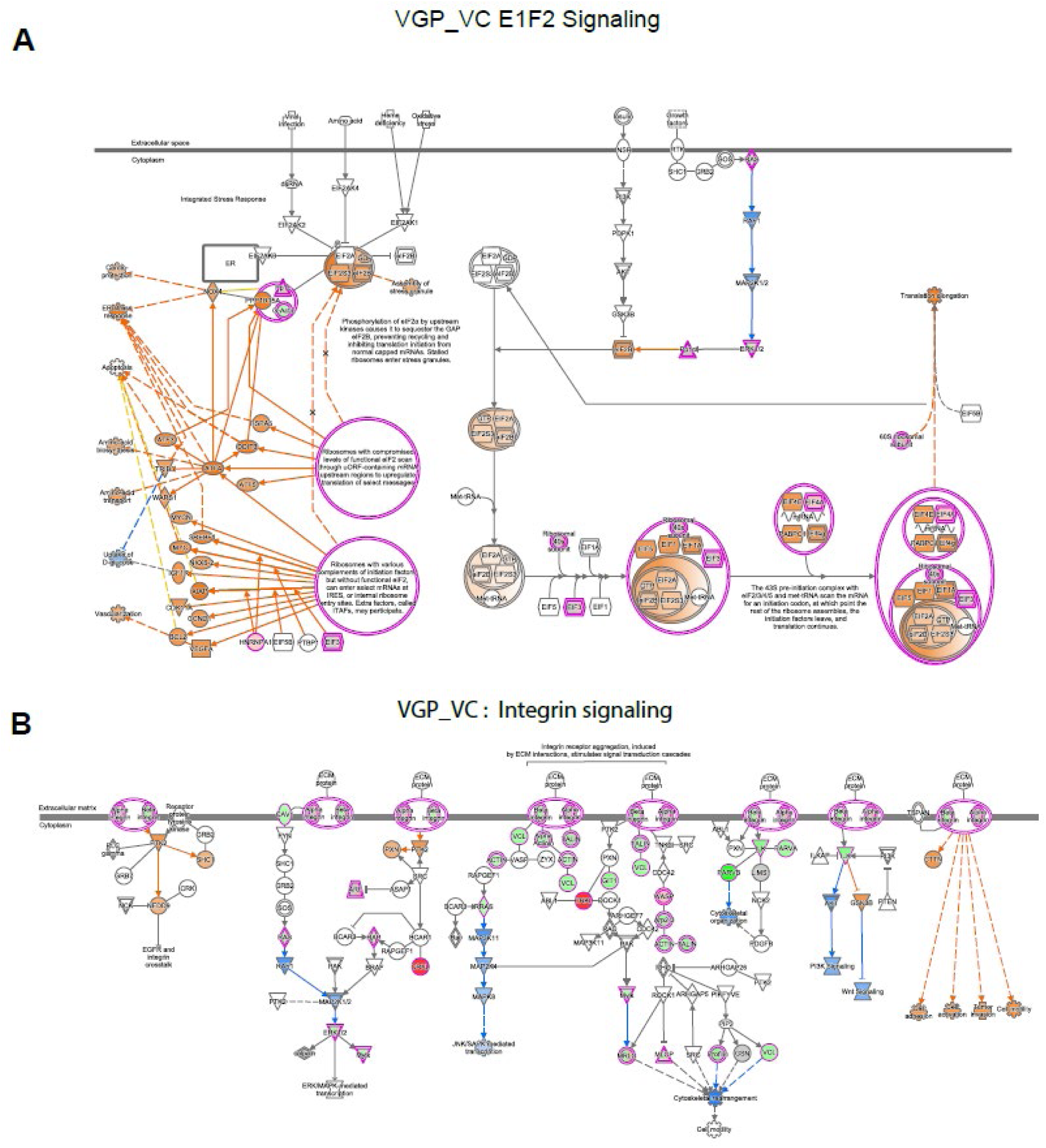
IPA canonical pathway maps highlighting EIF2 signaling and integrin signaling in functional vein grafts (VGP vs. VC). (A) EIF2 signaling. Ingenuity Pathway Analysis (IPA) canonical pathway map illustrating the EIF2 signaling cascade in the VGP vs. VC comparison. Upregulated components (highlighted in magenta) indicate activation of the integrated stress response, including increased EIF2 phosphorylation, ATF4-driven transcriptional programs, and downstream stress-adaptation modules. Predicted pathway activation is shown by orange arrows. (B) Integrin signaling. IPA canonical pathway map for integrin signaling in VGP vs. VC. Highlighted nodes (magenta) demonstrate upregulation of focal adhesion components, integrin receptor complexes, cytoskeletal regulators, and downstream MAPK/JNK-mediated signaling events. These findings are consistent with enhanced adhesion, mechanotransduction, and cytoskeletal remodeling in functional grafts.

**Figure S13.**
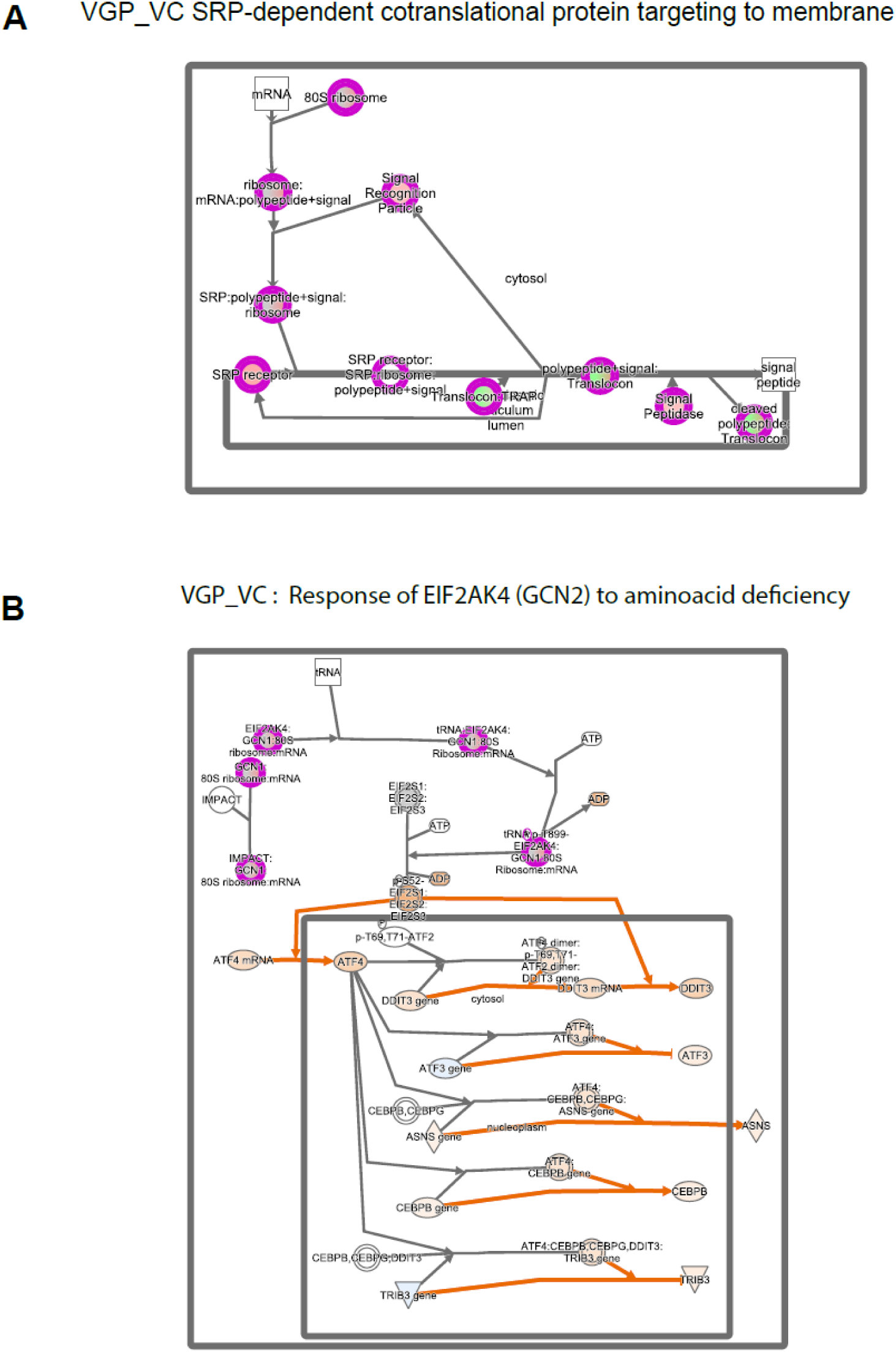
Canonical pathway maps illustrating translational and stress-response programs enriched in functional vein grafts (VGP vs. VC). (A) IPA canonical pathway diagram for the *SRP-dependent cotranslational protein targeting to the membrane*, highlighting upregulated components (magenta) in VGP relative to VC. This pathway represents increased engagement of ribosome–SRP complexes and endoplasmic reticulum translocation machinery, consistent with enhanced cotranslational processing in functional grafts. (B) IPA canonical pathway diagram for the *Response of EIF2AK4 (GCN2) to amino acid deficiency*, showing activation of GCN2-ATF4–dependent stress adaptation modules in VGP. Upregulated nodes (magenta) and predicted activation flow (orange arrows) indicate enhanced amino acid sensing, integrated stress response signaling, and downstream transcriptional activation. These pathway maps summarize the key translational and stress-adaptive mechanisms identified in the VGP vs. VC comparison.

**Figure S14.**
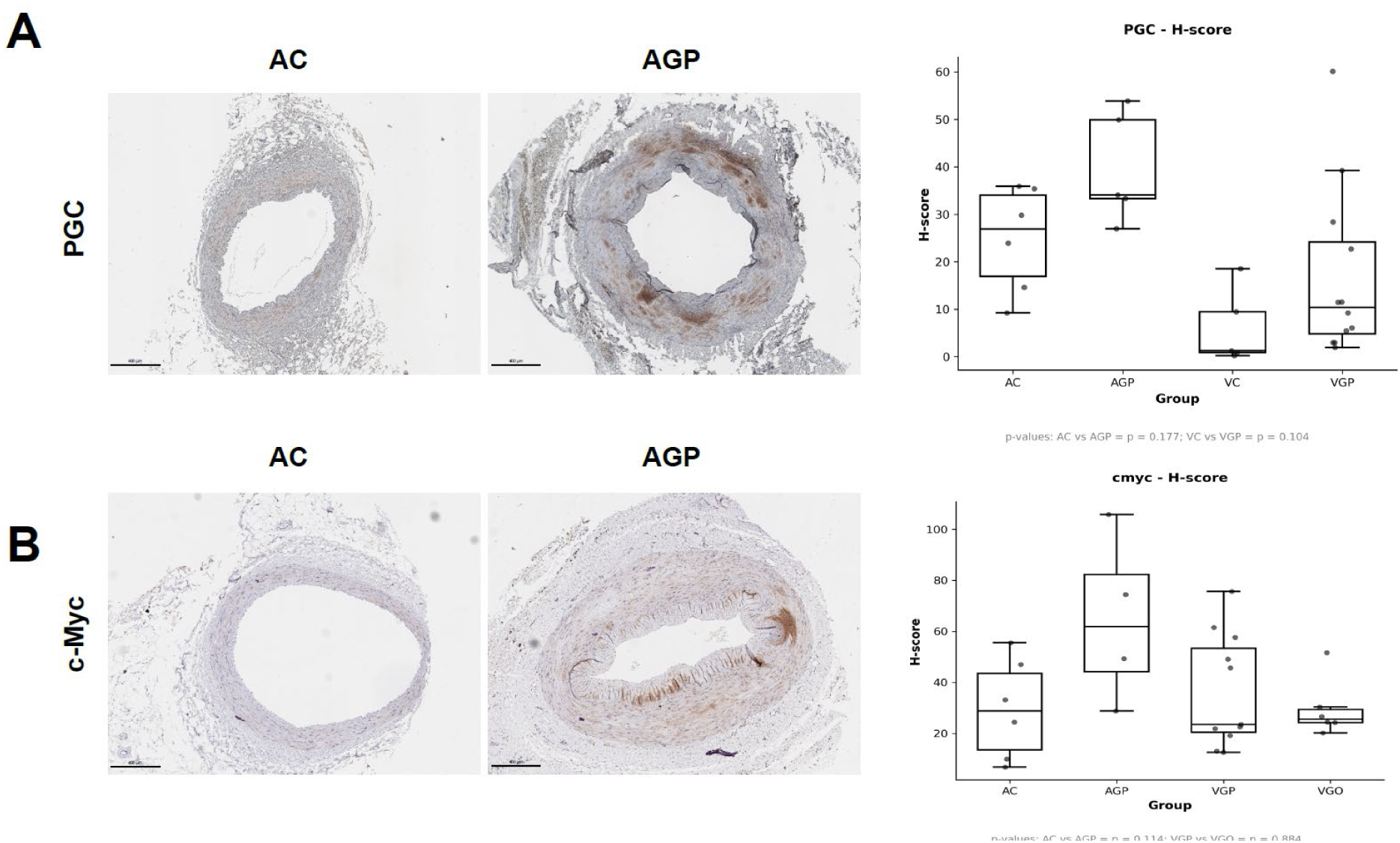
Immunohistochemical assessment of PGC-1α and c-Myc expression across graft conditions. (A) Representative immunohistochemical staining of PGC-1α in native arteries (AC) and patent arterial grafts (AGP) (left), with the corresponding H-score distribution across groups (right). PGC-1α showed an upward trend in both arterial and venous grafts (AGP > AC and VGP > VC), although the differences did not reach statistical significance (AC vs. AGP, P=0.177; VC vs. VGP, P=0.104). (B) Representative staining and H-score comparison for c-Myc. c-Myc expression was higher in AGP than in AC, but the difference was not statistically significant (P=0.114). No significant differences were observed between VGP and VGO (P=0.884). Scale bars are shown on each image

---

**Supplementary Table 1.**
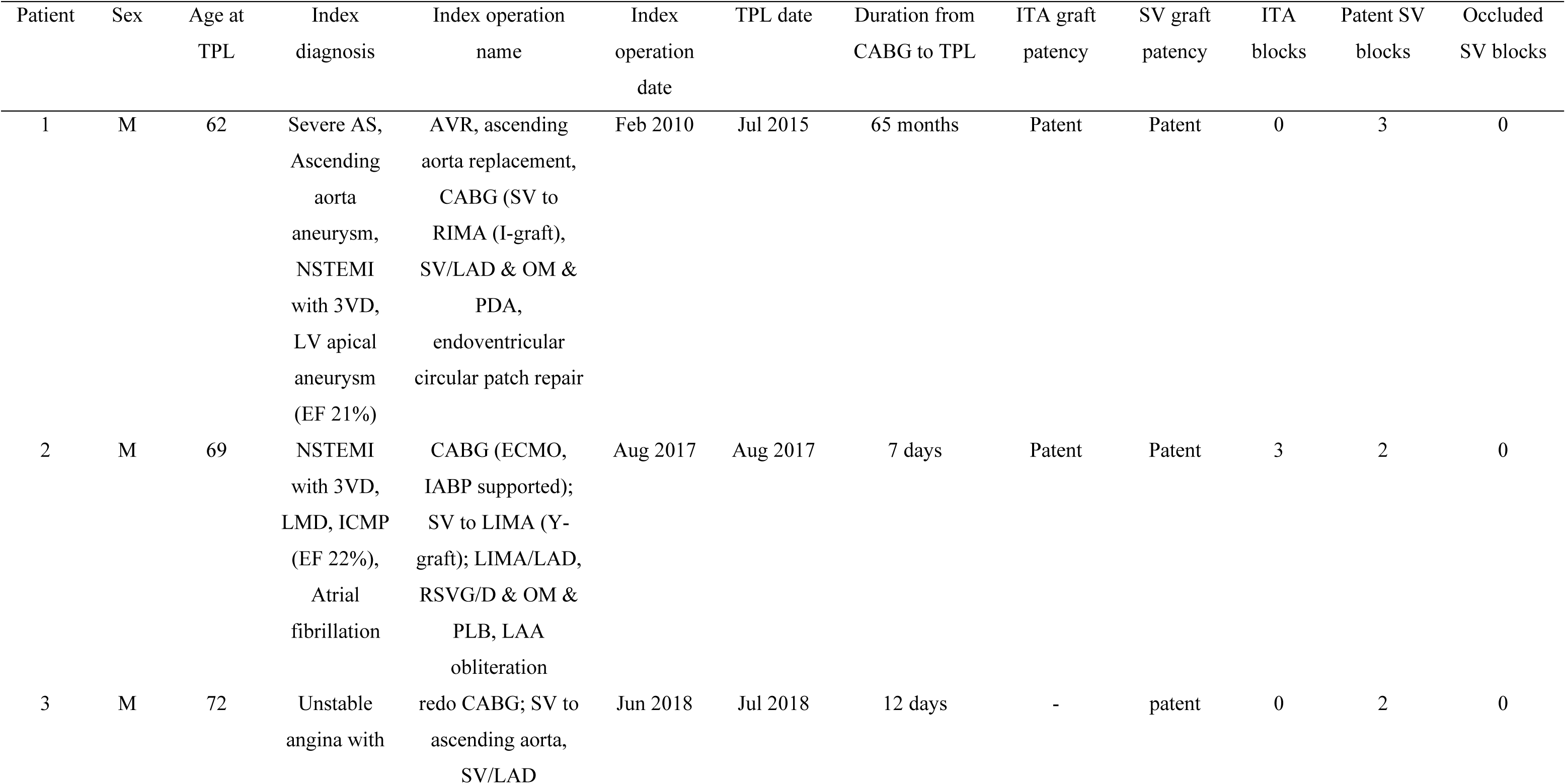
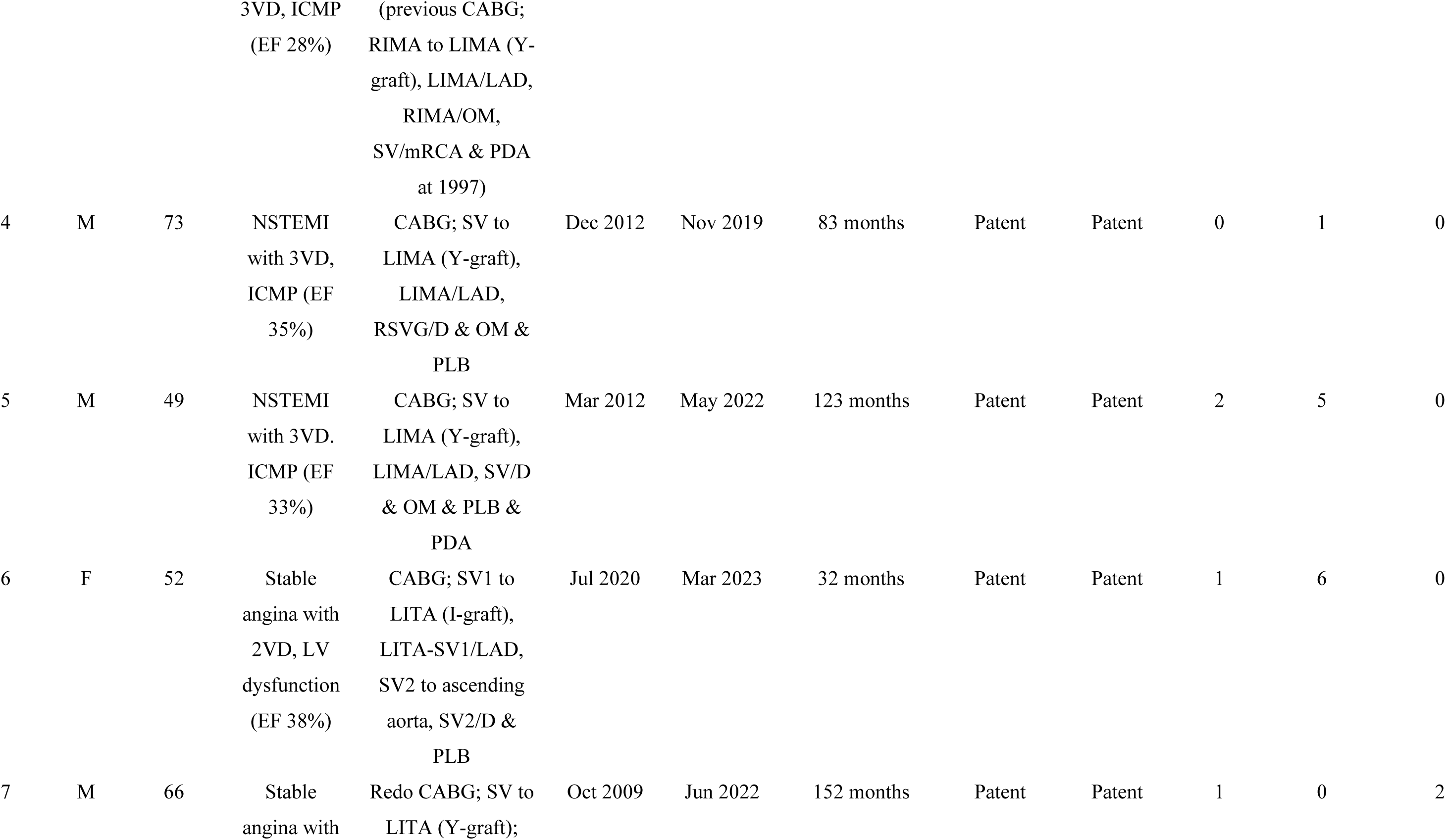
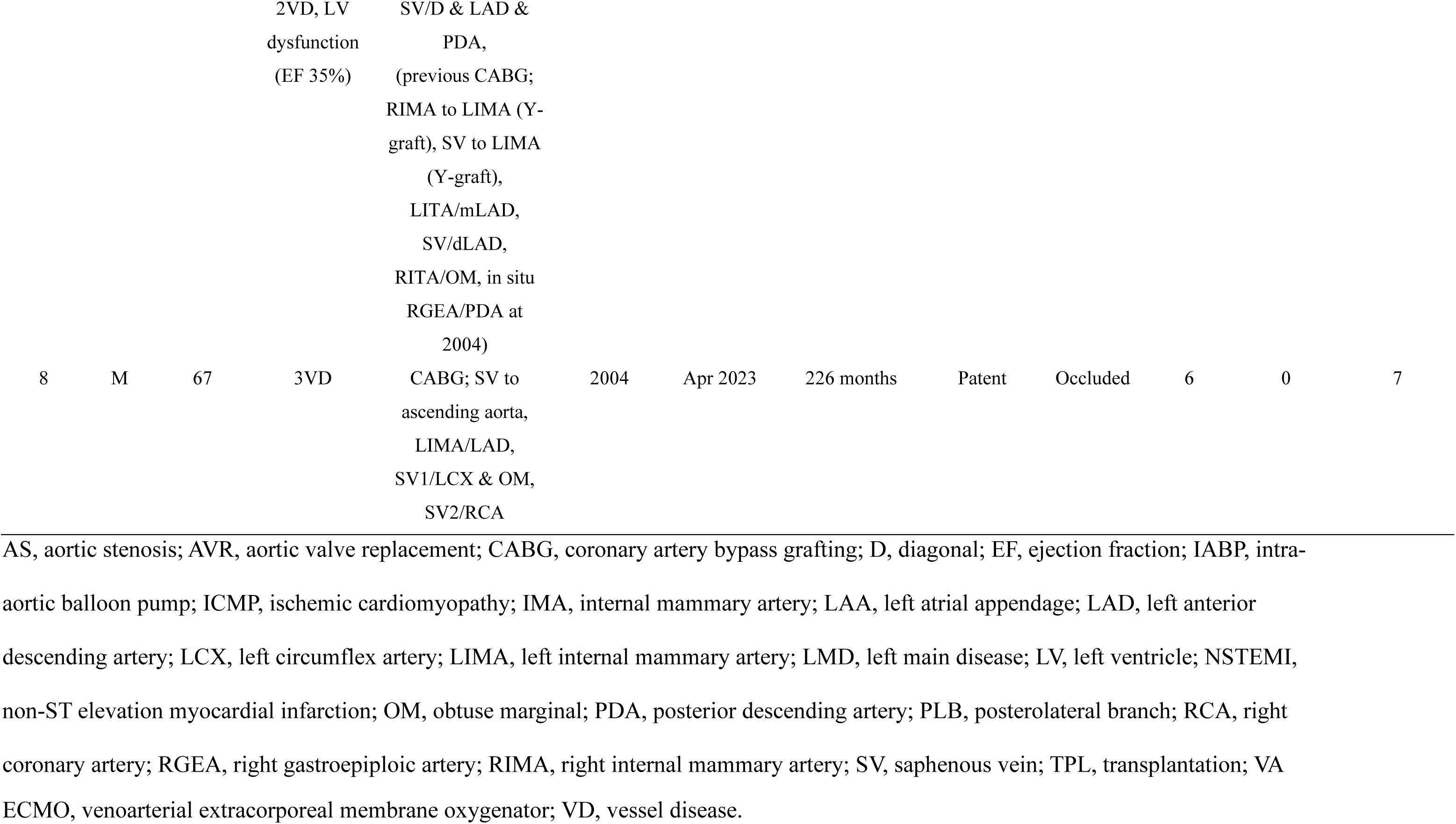
Patient characteristics.

| Patient | Sex | Age at<br>TPL | Index<br>diagnosis | Index operation<br>name | Index<br>operation<br>date | TPL date | Duration from<br>CABG to TPL | ITA graft<br>patency | SV graft<br>patency | ITA<br>blocks | Patent SV<br>blocks | Occluded<br>SV blocks |
| --- | --- | --- | --- | --- | --- | --- | --- | --- | --- | --- | --- | --- |
| 1 | M | 62 | Severe AS,<br>Ascending<br>aorta<br>aneurysm,<br>NSTEMI<br>with 3VD,<br>LV apical<br>aneurysm<br>(EF 21%) | AVR, ascending<br>aorta replacement,<br>CABG (SV to<br>RIMA (I-graft),<br>SV/LAD & OM &<br>PDA,<br>endoventricular<br>circular patch repair | Feb 2010 | Jul 2015 | 65 months | Patent | Patent | 0 | 3 | 0 |
| 2 | M | 69 | NSTEMI<br>with 3VD,<br>LMD, ICMP<br>(EF 22%),<br>Atrial<br>fibrillation | CABG (ECMO,<br>IABP supported);<br>SV to LIMA (Y-graft); LIMA/LAD,<br>RSVG/D & OM &<br>PLB, LAA<br>obliteration | Aug 2017 | Aug 2017 | 7 days | Patent | Patent | 3 | 2 | 0 |
| 3 | M | 72 | Unstable<br>angina with | redo CABG; SV to<br>ascending aorta,<br>SV/LAD | Jun 2018 | Jul 2018 | 12 days | - | patent | 0 | 2 | 0 |
|  |  |  | 3VD, ICMP<br>(EF 28%) | (previous CABG;<br>RIMA to LIMA (Y-graft), LIMA/LAD,<br>RIMA/OM,<br>SV/mRCA & PDA<br>at 1997) |  |  |  |  |  |  |  |  |
| 4 | M | 73 | NSTEMI<br>with 3VD,<br>ICMP (EF<br>35%) | CABG; SV to<br>LIMA (Y-graft),<br>LIMA/LAD,<br>RSVG/D & OM &<br>PLB | Dec 2012 | Nov 2019 | 83 months | Patent | Patent | 0 | 1 | 0 |
| 5 | M | 49 | NSTEMI<br>with 3VD.<br>ICMP (EF<br>33%) | CABG; SV to<br>LIMA (Y-graft),<br>LIMA/LAD, SV/D<br>& OM & PLB &<br>PDA | Mar 2012 | May 2022 | 123 months | Patent | Patent | 2 | 5 | 0 |
| 6 | F | 52 | Stable<br>angina with<br>2VD, LV<br>dysfunction<br>(EF 38%) | CABG; SV1 to<br>LITA (I-graft),<br>LITA-SV1/LAD,<br>SV2 to ascending<br>aorta, SV2/D &<br>PLB | Jul 2020 | Mar 2023 | 32 months | Patent | Patent | 1 | 6 | 0 |
| 7 | M | 66 | Stable<br>angina with | Redo CABG; SV to<br>LITA (Y-graft); | Oct 2009 | Jun 2022 | 152 months | Patent | Patent | 1 | 0 | 2 |
|  |  |  | 2VD, LV<br>dysfunction<br>(EF 35%) | SV/D & LAD &<br>PDA,<br>(previous CABG;<br>RIMA to LIMA (Y-graft), SV to LIMA<br>(Y-graft),<br>LITA/mLAD,<br>SV/dLAD,<br>RITA/OM, in situ<br>RGEA/PDA at<br>2004) |  |  |  |  |  |  |  |  |
| 8 | M | 67 | 3VD | CABG; SV to<br>ascending aorta,<br>LIMA/LAD,<br>SV1/LCX & OM,<br>SV2/RCA | 2004 | Apr 2023 | 226 months | Patent | Occluded | 6 | 0 | 7 |

AS, aortic stenosis; AVR, aortic valve replacement; CABG, coronary artery bypass grafting; D, diagonal; EF, ejection fraction; IABP, intra-aortic balloon pump; ICMP, ischemic cardiomyopathy; IMA, internal mammary artery; LAA, left atrial appendage; LAD, left anterior descending artery; LCX, left circumflex artery; LIMA, left internal mammary artery; LMD, left main disease; LV, left ventricle; NSTEMI, non-ST elevation myocardial infarction; OM, obtuse marginal; PDA, posterior descending artery; PLB, posterolateral branch; RCA, right coronary artery; RGEA, right gastroepiploic artery; RIMA, right internal mammary artery; SV, saphenous vein; TPL, transplantation; VA
ECMO, venoarterial extracorporeal membrane oxygenator; VD, vessel disease.

**Supplementary Table 2.**
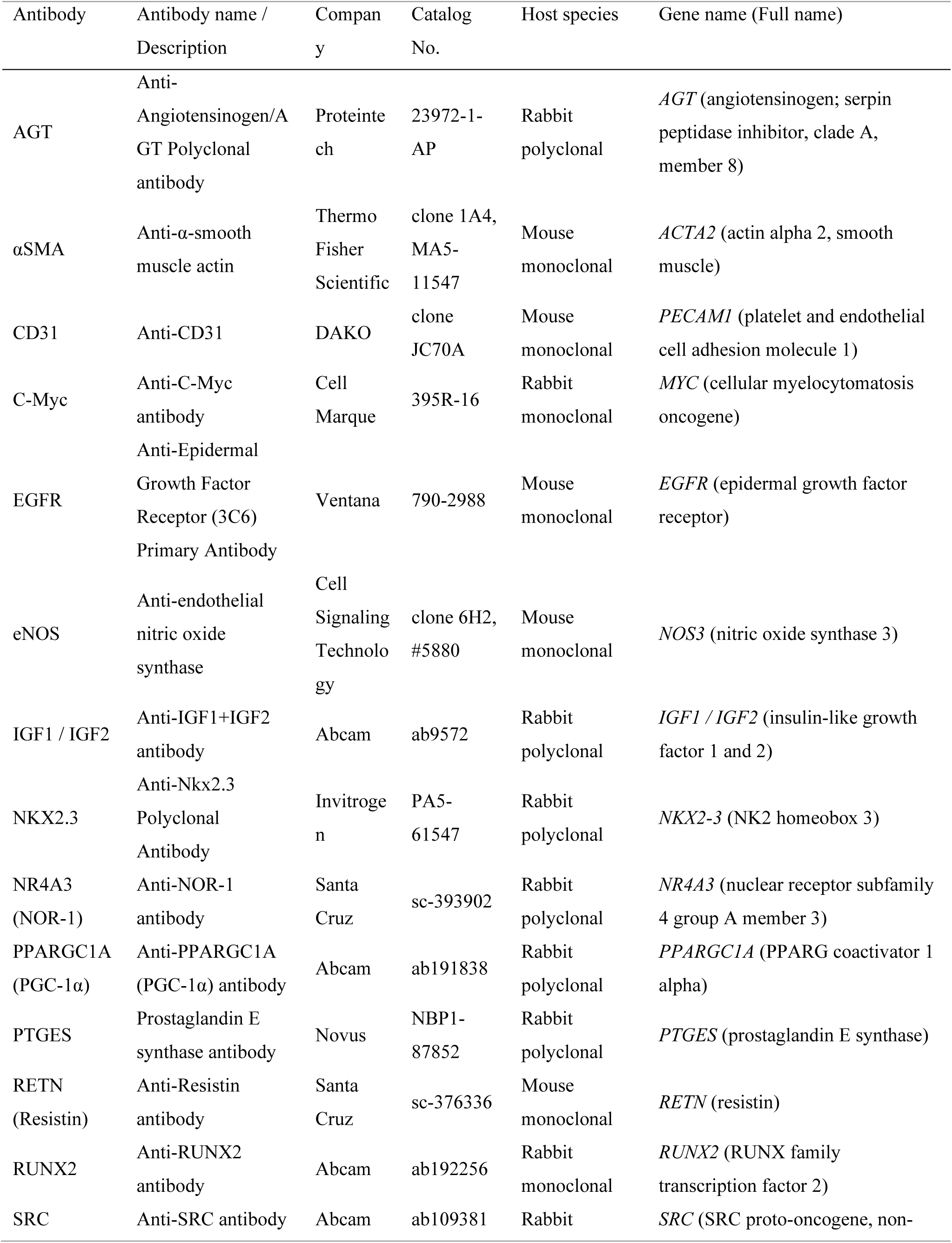
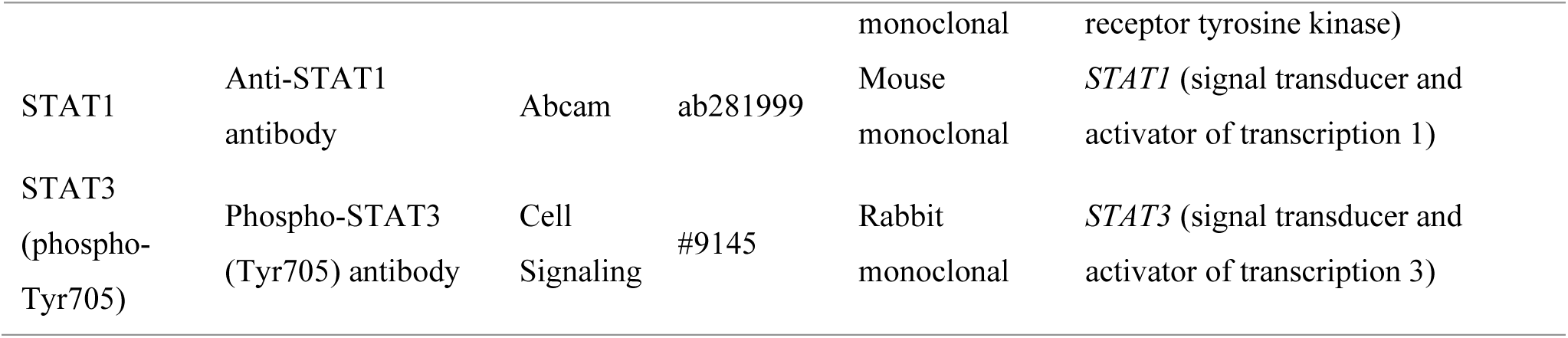
Antibodies used for immunohistochemistry.

